# Dynamic expression range expansion of ECF sigma factor-dependent synthetic regulons by promoter context engineering

**DOI:** 10.64898/2026.09.15.751809

**Authors:** Christian Rauch, Doreen Meier, Vinca Seiler, Eslam M. Elsayed, Anke Becker

**Affiliations:** Center for Synthetic Microbiology (SYNMIKRO), Philipps-Universität Marburg, Karl-von-Frisch-Straße 14, 35032 Marburg, Germany; Department of Biology, Philipps-Universität Marburg, Germany, Karl-von-Frisch-Straße 8, 35032 Marburg, Germany; Microbes-for-Climate (M4C) Cluster of Excellence, Philipps-Universität Marburg, Karl-von-Frisch-Straße 14, 35032 Marburg, Germany

## Abstract

Alternative σ factors enable bacteria to reprogram transcription in response to environmental cues and provide powerful tools for synthetic gene regulation. Extracytoplasmic function (ECF) σ factors have recently been implemented as orthogonal transcriptional switches in several bacteria, including the α-proteobacterium *Sinorhizobium meliloti*. Here, we investigated how promoter sequence context influences the activity and specificity of heterologous ECF-dependent core promoters. Combining ECF core promoters from *Pseudomonas syringae* and *Escherichia coli* with flanking sequences derived from strong promoters of *S. meliloti* and other proteobacteria increased promoter activity by up to 45-fold while preserving σ factor specificity. We identified a short upstream AₙT motif, resembling a minimal UP element, as a key determinant of promoter strength. Targeted mutagenesis confirmed its role in supporting transcription initiation. Engineering promoter-flanking sequences together with the AₙT motif generated promoter libraries spanning up to a 75-fold range of activities. Importantly, promoter activity hierarchies were maintained within synthetic multi-gene regulons controlled by a single ECF master regulator, demonstrating modular and predictable gene expression. Our results establish promoter environment engineering as a robust strategy for expanding the dynamic range of orthogonal ECF regulatory systems and facilitate the scalable design of synthetic transcriptional programs in bacteria.

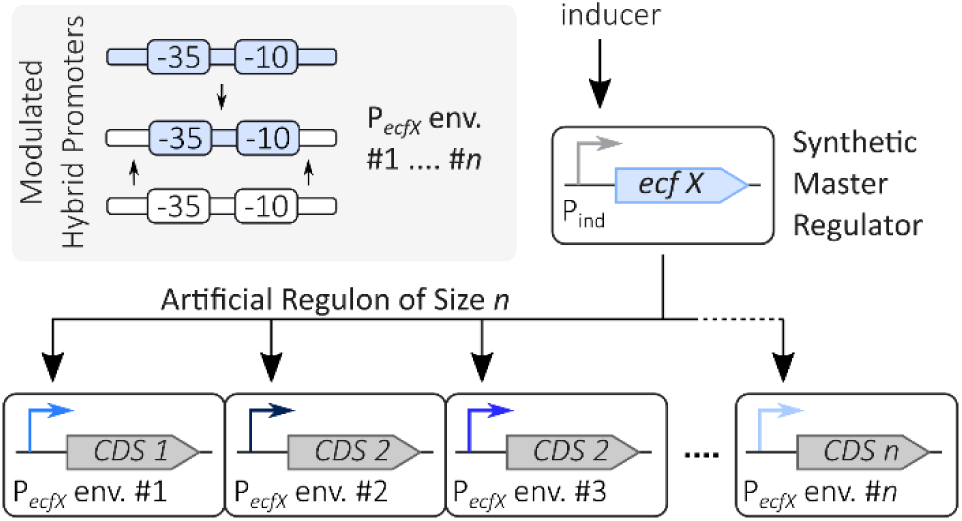

## Introduction

Microbial biotechnology increasingly leverages synthetic biology approaches to produce natural products and other valuable compounds in engineered bacterial hosts (Fang *et al*., 2017; Hug *et al*., 2020; Ray *et al*., 2022; Onyeaka and Ekwebelem, 2023; Malcı *et al*., 2024; Okoye *et al*., 2024; Yang *et al*., 2024; Boada *et al*., 2025; Gomes *et al*., 2025; Stegman *et al*., 2025). As synthetic pathways become more sophisticated, incorporating multiple transcription units and additional regulatory layers, precise and predictable control of gene expression becomes increasingly important (Fang *et al*., 2017; Boada *et al*., 2025). One promising strategy is the repurposing of bacterial signal transduction mechanisms as orthogonal regulatory modules.

Bacteria employ diverse regulatory systems to couple environmental and intracellular signals to transcriptional responses. In addition to transcription factor- and riboregulator-based regulation, transcriptional space is partitioned into distinct regulons controlled by alternative σ factors (Gruber and Gross, 2003). While housekeeping σ factors (RpoD or σ^70^ in *Escherichia coli*) direct transcription of genes involved in growth and central metabolism, alternative σ factors redirect the RNA polymerase (RNAP) to dedicated promoter classes and thereby enable specialized transcriptional programs. Among them, members of the σ^70^ family have attracted particular interest as engineering tools because they directly determine promoter recognition and can establish orthogonal transcriptional circuits.

Group IV σ factors, also known as extracytoplasmic function σ factors (ECFs), are especially attractive for synthetic biology applications. ECFs are compact regulators composed primarily of σ_2_ and σ_4_ domains that recognize the −10 and −35 promoter elements, respectively (Lonetto *et al*., 1994; Helmann, 2002; Staroń *et al*., 2009; Feklístov *et al*., 2014; Li *et al*., 2019; Lin *et al*., 2019). Many ECFs are negatively controlled by specific binding of anti-σ factors (Helmann, 1999, 2002). ECFs are widely distributed throughout the bacterial kingdom and exhibit extensive diversity in promoter specificity, regulatory mechanisms and regulon size (Staroń *et al*., 2009; Todor *et al*., 2020; Casas-Pastor *et al*., 2021). This diversity has enabled the development of synthetic transcriptional switches based on heterologous ECFs and cognate promoters (ECF switches) in multiple bacterial species and *in vitro* (Chen and Arkin, 2012; Shin and Noireaux, 2012; Rhodius *et al*., 2013; Zong *et al*., 2017; Bervoets *et al*., 2018; Pinto *et al*., 2018, 2019; Van Brempt *et al*., 2020; Zhao *et al*., 2022; Meier *et al*., 2024; Patel *et al*., 2024; Boada *et al*., 2025; Wolters *et al*., 2026).

Synthetic regulons are particularly attractive for the genetic implementation of synthetic metabolic modules, as they enable efficient tuning through manipulation of a single regulatory node and minimize regulatory burden. A key requirement for constructing synthetic ECF regulons is the availability of promoter sets that combine high σ-factor specificity with tunable and predictable expression strengths. Previous studies generated promoter libraries by introducing mutations within the spacer or the entire core region (defined as the −35 box/spacer/-10 box region) of ECF- and group III σ factor-dependent promoters, producing variants that mediated broad expression ranges in *Escherichia coli* and *Streptomyces venezuelae* (Zong *et al*., 2017; Bervoets *et al*., 2018; Zhao *et al*., 2022). Such a library has enabled high-throughput optimization of a four-gene heterologous biosynthetic pathway by high-throughput screening of operon configurations (Van Brempt *et al*., 2022), and machine learning-guided exploration of expression spaces of synthetic pathway modules has been demonstrated to significantly accelerate metabolic engineering (Van Brempt *et al*., 2022; Schulz-Mirbach *et al*., 2026). This highlights the potential of synthetic ECF regulons for rational genetic implementation of *in silico* optimized pathway variants.

However, varying promoter activity through mutations within the core region may alter its specificity and has little potential for enhancing overall promoter strength. Engineering promoter-flanking sequences represents an attractive alternative for promoter tuning since these regions contribute to RNAP-promoter interactions. Structural and functional studies have shown that regions outside the −35 and −10 elements can substantially influence transcription initiation by RNAP-ECF holoenzymes. Interactions between RNAP and upstream promoter DNA can enhance transcription through UP elements in both RpoD-dependent and ECF-dependent promoters (Estrem *et al*., 1998, 1999; Rhodius *et al*., 2012, 2013; Zong *et al*., 2017; Presnell *et al*., 2019; Todor *et al*., 2020). UP elements are typically AT-rich motifs composed of a proximal and distal site, each of which can individually bind the carboxy-terminal domain of an RNAP α subunit (α-CTD) and stimulate transcription (Estrem *et al*., 1998, 1999). Beyond upstream sequences, downstream promoter regions can also affect transcriptional output, for example through the core recognition element (CRE) (Lin *et al*., 2019). Despite these findings, the potential of promoter-flanking sequences as engineering targets for tuning ECF-dependent gene expression has received little attention.

We investigated the effect of promoter-flanking sequences on ECF-dependent promoters in *Sinorhizobium meliloti*, an alphaproteobacterial model for nitrogen-fixing root nodule symbiosis and an emerging platform for biotechnological applications (Galibert *et al*., 2001; Jones *et al*., 2007; Dong *et al*., 2016). Previously, we established heterologous ECF switches in *S. meliloti* (Meier *et al*., 2024). Here, we show that engineering the sequence environment surrounding ECF-dependent core promoters enables extensive modulation of promoter activity while preserving promoter specificity. By combining heterologous ECF core promoters with promoter environments derived from sequences native to either *S. meliloti* or other proteobacteria, and by mutating upstream sequence elements, we generated libraries of ECF-specific promoter variants whose transcriptional activities varied by up to 75-fold. We identify a short UP element-like A_n_T motif as a major determinant of promoter strength, and demonstrate that the various promoter strengths measured in single-promoter reporter assays are maintained within synthetic multi-gene regulons controlled by a single ECF master regulator. These findings promote promoter-environment engineering as a powerful strategy for constructing predictable and tunable ECF-dependent transcriptional programs and facilitate the implementation of complex synthetic pathway modules in *Sinorhizobium* and potentially other bacterial hosts.

## Results

### A salicylate-inducible promoter improves resolution of both strong and weak ECF switches in *S. meliloti*

To test the influence of the surrounding sequence context on the activity of heterologous ECF core promoters (−35 box/spacer/-10 box) in *S. meliloti*, an assay system consisting of two compatible, mobilizable single-copy plasmids was established. The system consisted of (i) an input module comprising the respective *ecf* gene controlled by an inducible promoter and (ii) an output module comprising the *luxCDABE* luminescence reporter operon controlled by a promoter (P*_ecf_*) depending on the input module-encoded ECF (Fig. 1A). The applicability of this system for our studies into the influence of ECF promoter sequence environments relies strongly on the properties of the inducible promoter controlling *ecf* gene expression. Ideally, a promoter that exhibits barely detectable basal activity in the off-state, strong gene expression in the on-state, and linear dependence of gene expression on inducer concentration is used. However, previously characterized inducible promoters did not fully meet these requirements in *S. meliloti*.

**Fig. 1:**
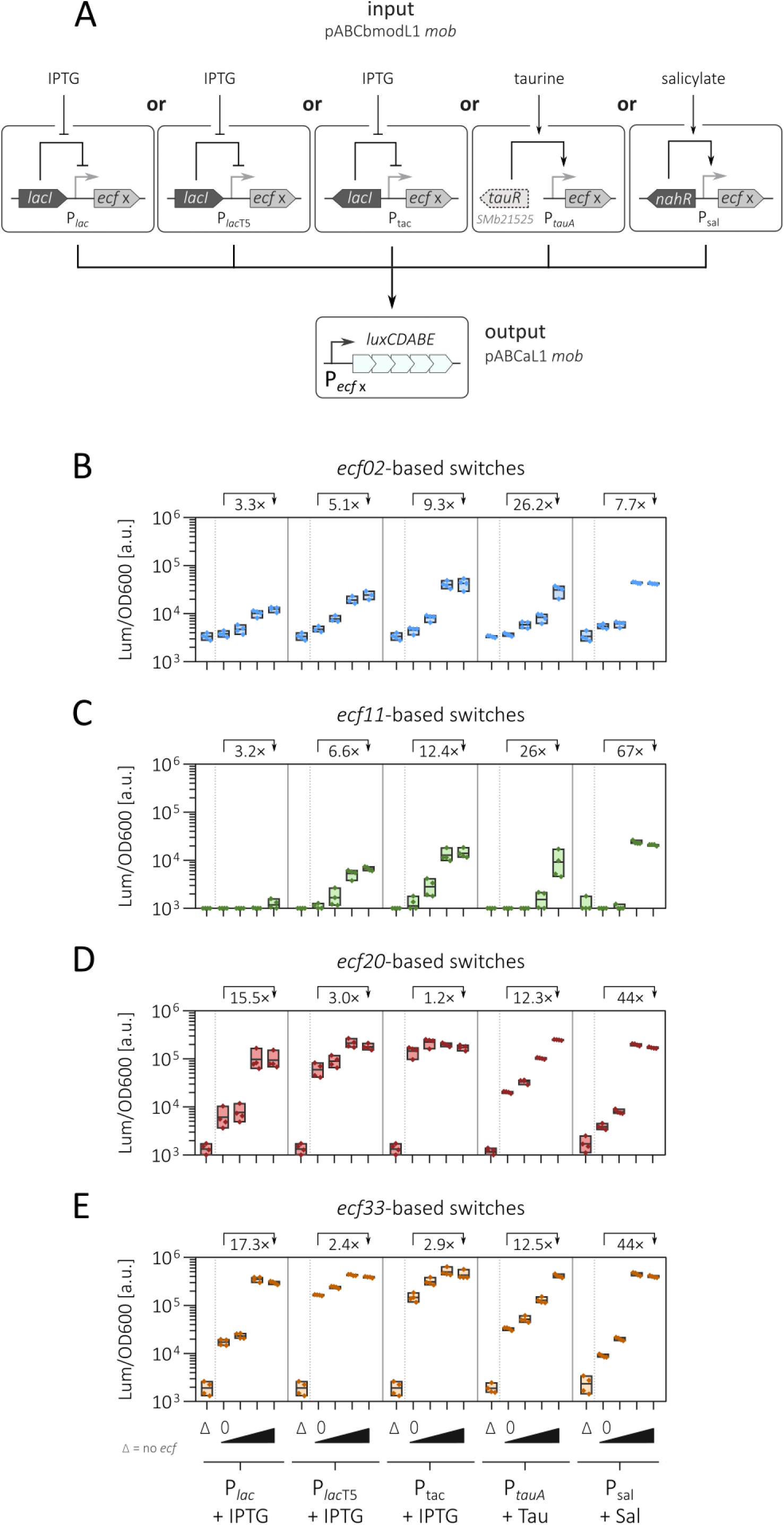
Characterization of ECF switches in *S. meliloti* utilizing diverse inducible input promoters. **(A)** Schematic representation of two-plasmid-borne ECF switches. The input module on the single copy plasmid pABCbmodL1 *mob* consisted of different *ecf* genes transcriptionally controlled either by derivatives of the IPTG-inducible *E. coli lacI*-P*_lac_* regulator/promoter pair, the taurine-inducible P*_tauA_* regulated by the endogenous GntR-family activator TauR (encoded by *SMb21525* on the endogenous chromid pSymB), or the salicylate-inducible *nahR*/P_sal_ regulator/promoter pair (Mostafavi *et al*., 2014; Meyer *et al*., 2019; Meier *et al*., 2024; Faber *et al*., 2025). The *ecf* gene was preceded by a standardized 5’-UTR (Meier *et al*., 2024). The output module of the single-copy plasmid pABCaL1 *mob* consisted of a P*_ecf_*promoter, which was cognate to the ECF encoded by the input module. This promoter was transcriptionally fused to a *luxCDABE* reporter cassette, which was preceded by a standardized 5’-UTR. The system was designed to enable flexible incorporation of the inducible promoter/transcription regulator gene pair as well as P*_ecf_* and the *ecf* gene using Golden Gate cloning. **(B-E)** Luminescence output of *ecf02*-**(B)**, *ecf11*-**(C)**, *ecf20*-**(D)**, and *ecf33*-based **(E)** two-plasmid switches in *S. meliloti* Δ*ecf*/Δ*anti-σ* in MOPS-buffered minimal medium 9 h after the addition of the respective inducer. Strains were induced with different concentrations of IPTG (0 µM, 10 µM, 50 µM, 500 µM), taurine (Tau; 0 mM, 0.2 mM, 1 mM, 3 mM), or salicylic acid (Sal; 0 µM, 5 µM, 25 µM, 50 µM). Strains lacking the *ecf* gene on the input module plasmid are designated as “Δ”. Each strain was measured in four biological replicates. Numerical values represent the average fold induction between the uninduced off-state and fully induced on-state.

Therefore, we initially characterized various inducible promoters in *S. meliloti* and subsequently re-evaluated a subset of previously characterized ECF switches (Meier *et al*., 2024) using the most promising promoter candidates. We generated single-copy reporter plasmids carrying the luminescence reporter cassette (*luxCDABE*) controlled by nine different transcription regulator gene/inducible promoter pairs (Mostafavi *et al*., 2014; Meyer *et al*., 2019; Meier *et al*., 2024; Faber *et al*., 2025). Basal and fully induced promoter activities were measured in *S. meliloti* Δ*ecf*/Δ*anti-σ*, a strain lacking the 11 endogenous ECFs and associated anti-σ factors (Lang *et al*., 2018) (Fig. S1A-B). This strain was later also used for all ECF switch characterizations to exclude cross-talk. Five of these inducible promoters were found to be functional, namely P*_lac_*, P*_lac_*_T5_, P_tac_, P*_tauA_*, and P_sal_ (Fig. S1C), and the best performing promoters P*_sal_*and P*_tauA_* showed a linear dose-response behavior in the inducer concentration range of 0.05 to 3 mM taurine and 2.5 to 50 µM salicylate, respectively (Fig. S1D, E).

Therefore, P*_sal_* and P*_tauA_* as well as for comparison, P*_lac_*, P*_lac_*_T5_, and P*_tac_* were incorporated into our two-plasmid ECF switch test system. For further evaluation we chose four heterologous ECF/promoter pairs that previously exhibited strong (ECF20/P*_ecf20_* from *Pseudomonas fluorescens*, and ECF33/P*_ecf33_* from *Rhodospeudomonas palustris* and *Bradyrhizobium japonicum* for ECF and promoter, respectively), medium (ECF02/P*_ecf02_* from *E. coli*), or weak (ECF11/P*_ecf11_*from *Vibrio parahaemolyticus* and *Pseudomonas syringae* for ECF and promoter, respectively) activities and minimal cross-talk in *S. meliloti* (Meier *et al*., 2024). Complete sequences of all ECFs and their cognate promoters are presented in Tables S1 and S2, respectively. OD600 (optical density at 600 nm) and luminescence (Lum) of cultures of *S. meliloti* test strains carrying the resulting input and output plasmids as well as of associated control strains lacking the *ecf* gene in the input plasmid were measured nine hours after addition of the respective inducers (Fig. 1B-E).

Overall, the salicylate-inducible system demonstrated optimal regulatory properties: strong ECF switches (ECF20/P*_ecf20_*, ECF33/P*_ecf33_*) achieved the highest fold-change through a low basal activity in the off-state. Leaky expression from the P*_tauA_* promoter was 5.2- and 3.7-fold for ECF20/P*_ecf20_* and ECF33/P*_ecf33_*, respectively, compared to the P_sal_ system. While this effect was less pronounced for the weak ECF11/P*_ecf11_*and the medium ECF02/P*_ecf02_* switches, both of them showed an increased on-state activity of 2.3- and 1.4-fold, respectively, with P_sal_ as input promoter compared to P*_tauA_*. This suggests that the *nahR*/P_sal_ regulator-promoter pair is most suitable for induction of ECF switches across all activity levels in *S. meliloti.* We therefore chose this inducible promoter for all following characterizations of ECF switches in this study.

### Promoter environments modulate ECF promoter activity and preserve promoter specificity

We chose the ECF/promoter pairs ECF02/P*_ecf02_* and ECF11/P*_ecf11_* to test for the influence of promoter sequence environments on activity and specificity of these heterologous ECF switches. We focused on these ECF switches as they exhibited medium and weak activity, respectively, allowing for identifying variants with increased as well as reduced activity levels. We defined the heterologous ECF core promoter (27 to 34 bp) as the −35 and −10 boxes, and the spacer connecting both, with the two boxes constituting the specific ECF binding motifs. The core promoters were equipped with upstream (27 to 33 bp) and downstream (29 to 30 bp) sequences of various, well-established donor promoters, to generate hybrid promoter regions (Fig. 2A). A detailed depiction of this hybrid design is provided in Fig. S2.

**Fig. 2:**
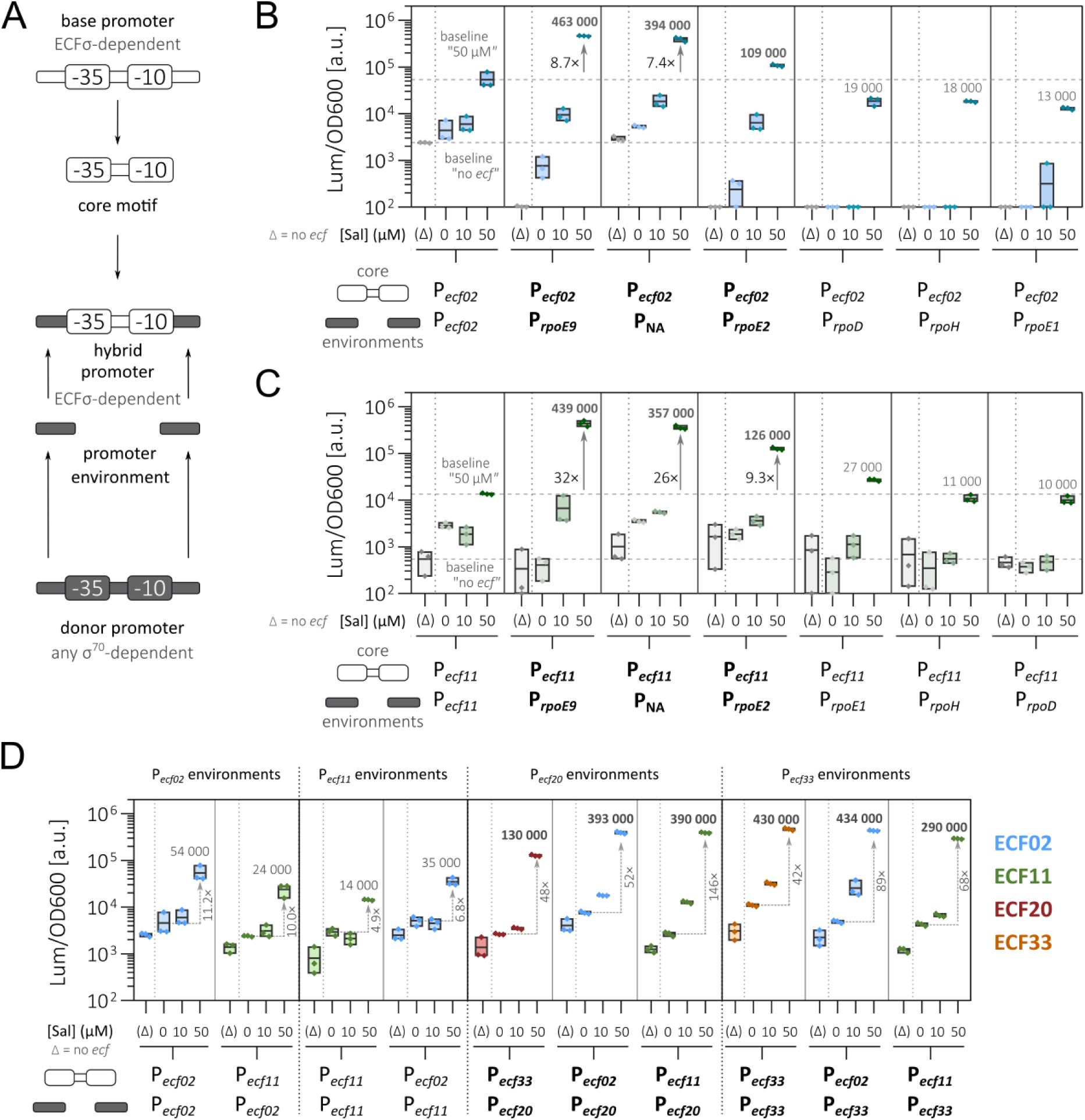
Characterization of hybrid ECF-dependent promoters in *S. meliloti*. **(A)** Schematic representation of the procedure employed to engineer hybrid ECF promoters. The ECF-dependent core promoter (light grey) and the donor promoter controlled by σ factors belonging to various groups of the σ^70^ family (dark grey) were aligned with respect to their TSS (+1). The core region of the base promoter (spanning from the −35 box to the −10 box, including the spacer sequence) was fused with environmental sequences from the donor promoter (see Fig. S2 for details). **(B, C)** Activities of ECF02-(B) and ECF11*-*dependent (C) switches with original and hybrid promoters, the latter containing P*_ecf02_* or P*_ecf11_* core regions and sequence environments derived from various σ^70^-family σ factor-dependent *S. meliloti* promoters (Schlüter *et al*., 2013). Grey numbers indicate average activity of fully induced strains. Black numbers indicate the average fold increase of the on-state of a given hybrid promoter compared to the respective wild type promoter. **(D)** Activities of ECF switches with original and hybrid promoters containing environments derived from heterologous ECF promoters. For the construction of hybrid promoters, core regions of P*_ecf02_* and P*_ecf11_* and environments derived from P*_ecf20_* and P*_ecf33_*, and either P*_ecf02_* or P*_ecf11_* were used. Promoters that exhibited strong activity (> 10^5^ Lum/OD600) are highlighted in bold. All ECF switches (B-D) were established in *S. meliloti* Δ*ecf*/Δ*anti-σ* using the two-plasmid system depicted in Fig. 1A. Each P*_ecf_* was paired with its cognate *ecf* according to the core region of the base promoter or as control with an empty input plasmid lacking the *ecf* gene (designated “(Δ)”). Strains were cultured in MOPS-buffered minimal medium in the presence of the indicated salicylate inducer concentrations. OD600 and luminescence were captured 9 h post-induction with three biological replicates per strain. Grey horizontal numbers indicate average activity of fully induced strains. Grey vertical numbers alongside arrows indicate the average fold change between the on- and the off-state of each switch.

The donor promoter environments were derived from the endogenous *S. meliloti* promoters indicated in Fig. 2B and C. Environments were derived from promoters recognized by the housekeeping σ factor RpoD (corresponding gene: *SMc03990*), the heat shock σ factors RpoH1/2 (*SMc02575*), and the endogenous ECFs RpoE1 (*SMc01418*), RpoE2 (*SMc01615*), and RpoE9 (*SMb20029*). Additionally, one donor promoter was included that was <u>n</u>ot <u>a</u>ssigned to a specific sigma factor (“NA”, corresponding gene: *SMc01180*) (Schlüter *et al*., 2013). Comparison with publicly available databases revealed no binding sites for transcription factors in the environments of these *S. meliloti* donor promoters (Taboada-Castro *et al*., 2024), indicating that these do not underly additional transcriptional regulation.

Furthermore, our study included the native environments from P*_ecf20_*and P*_ecf33_* (promoters of highly active heterologous ECF switches) as well as from P*_ecf02_* and P*_ecf11_* (promoters of heterologous ECF switches with medium and weak activities, respectively) (Fig. 2D). All promoter environments were extended 5 bp both upstream and downstream compared to previous studies (Rhodius *et al*., 2013; Meier *et al*., 2024) resulting in a total length of 90 bp per ECF-promoter (−65 to +25 relative to the TSS) to accommodate for full UP elements (Rhodius *et al*., 2012) and discriminator regions (Schlüter *et al*., 2013) potentially present in the donor promoters (Fig. S2; Table S2).

The hybrid promoters were tested for their activity with the ECF cognate to the core promoter using the two-plasmid assay system in *S. meliloti* described above. Test strains and control strains lacking the *ecf* gene were cultured with varying salicylate concentrations, and OD600 and luminescence were determined after nine hours (Fig. 2B-D). Notably, the engineered promoter environments had comparable effects on the performance of the ECF02- and ECF11-based switches. The P*_rpoE9_* promoter environment produced the highest luminescence activities, which were similar for full induction of *ecf02* and *ecf11* expression – (463 ± 8) × 10^3^ and (440 ± 50) × 10^3^ Lum/OD600, respectively (Fig. 2B, C). ECF switches with a lower on-state level showed a higher degree of variance in the activities of ECF02 and ECF11 core promoters flanked by the same promoter environments. For example, the core promoters with the P*_rpoE1_* environment resulted in (13.0 ± 0.7) × 10^3^ and (27.1 ± 1.3) × 10^3^ Lum/OD600 in the ECF02- and ECF11-dependent switches, respectively, corresponding to a 2.1-fold difference between both switches.

Changes in the environment of the core promoters also influenced promoter recognition by endogenous σ factors of the *S. meliloti* Δ*ecf*/Δ*anti-σ* test strain background, possessing the housekeeping σ factor RpoD, as well as the alternative non-ECF σ factors RpoH1, RpoH2, and RpoN. In the absence of the *ecf02* gene, most P*_ecf02_* promoter variants showed a decreased background activity (0.1 × 10^3^ Lum/OD600 or lower) as compared to the original P*_ecf02_*, which showed a high basal activity of approximately 2.5 × 10^3^ Lum/OD600 (Fig. 2B). The decrease in ECF-independent transcriptional activation of ECF-recognized promoters was inconsistent between P*_ecf02_* and P*_ecf11_* promoters. This suggests that the reduction of non-specific promoter activation resulted from the interplay of the core promoter and its sequence environment.

To assess whether the ECF11 and ECF02-specific core promoters maintained their specificities for their cognate ECFs despite being equipped with non-cognate P*_ecf_*-derived sequence environments (P*_ecf20_*, P*_ecf33_*; P*_ecf02_* or P*_ecf11_*), we tested their activation by eight previously characterized heterologous, non-cognate ECFs (Meier *et al*., 2024). This set included the four ECFs that recognize the donor promoters for the sequence environments. Each of the hybrid promoter-*luxCDABE* output variants were combined with the different P_sal_-*ecf* input variants using the established two-plasmid assay system in *S. meliloti*. The resulting strains were cultured and OD600 and luminescence were measured nine hours post-induction. We compared the on-state of each ECF/P*_ecf_* pair-containing strain to a control strain carrying the output module but lacking the *ecf* in the input module (Fig. 3). Within the set of nine ECFs, no unexpected cross-talk between hybrid promoters and non-cognate ECFs was observed. This suggests that the exchange of the promoter environments modulated promoter activity while maintaining specificity.

**Fig. 3:**
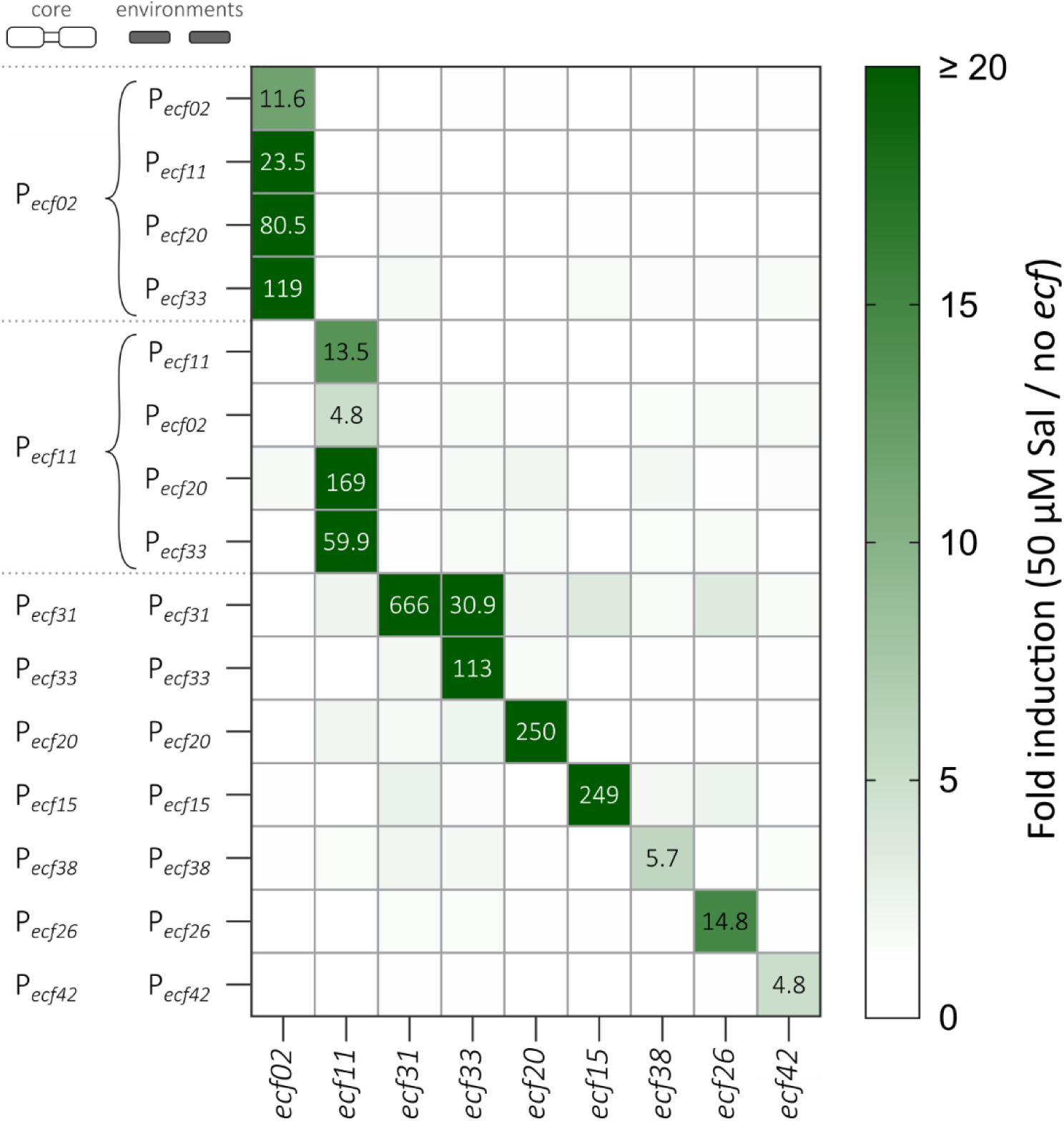
Orthogonality matrix of hybrid promoters among previously established other heterologous ECFs. Each cell represents an *S. meliloti* Δ*ecf*/Δ*anti-σ* strain carrying a two plasmid-based ECF switch whose activity in the on-state was compared to its respective “no *ecf*” control strain carrying the same output module (P*_ecf_*-*luxCDABE*). In the heat map, columns represent the respective *ecf* utilized in the input module of the switch while each row represents the P*_ecf_*-*luxCDABE* output module. On the left, core base promoters and environment-donating promoters for the (hybrid) promoters of the output module are indicated. Each strain was measured in at least three biological replicates. Strains were cultured in MOPS-buffered minimal medium and data was captured 9 h after addition of 50 µM salicylate.

### A short UP element-like A_n_T motif predicts highly active promoter environments

Promoter environments that conferred strong ECF switch activity (above 10^5^ Lum/OD600 in the on-state) did so with remarkable consistency, regardless of the core promoter and the associated ECF. To identify sequence features that might account for enhanced transcription initiation rates of strong hybrid promoters, we searched for AT-rich sequence stretches in the upstream promoter environments, since one prominent feature of bacterial promoters is the UP element (Ross *et al*., 1993; Estrem *et al*., 1998).

We identified a short motif consisting of multiple adenines followed by at least one thymine, here termed the A_n_T motif, which was present in the strong but absent from the weak hybrid promoters (Fig. 4A). When present in ECF-dependent promoters, the A_n_T motif was located 9 to 21 bp (for ECF02-dependent promoters) or 8 to 20 bp (for ECF11-dependent promoters) upstream of the 5’ end of −35 box and overlapped the region corresponding to the distal contact site of the RNAP α-CTD involved in UP element recognition. Since the C-terminal domain of the α-subunits preferentially interacts with narrow DNA minor grooves during UP element binding (Lara-Gonzalez *et al*., 2020), we calculated the minor groove width (MGW) of strong (Fig. 4B) and weak (Fig. 4C) upstream promoter environments. Promoters containing the A_n_T motif exhibited a reduced MGW of ≤ 3.5 Å, a characteristic feature of UP elements.

**Fig. 4:**
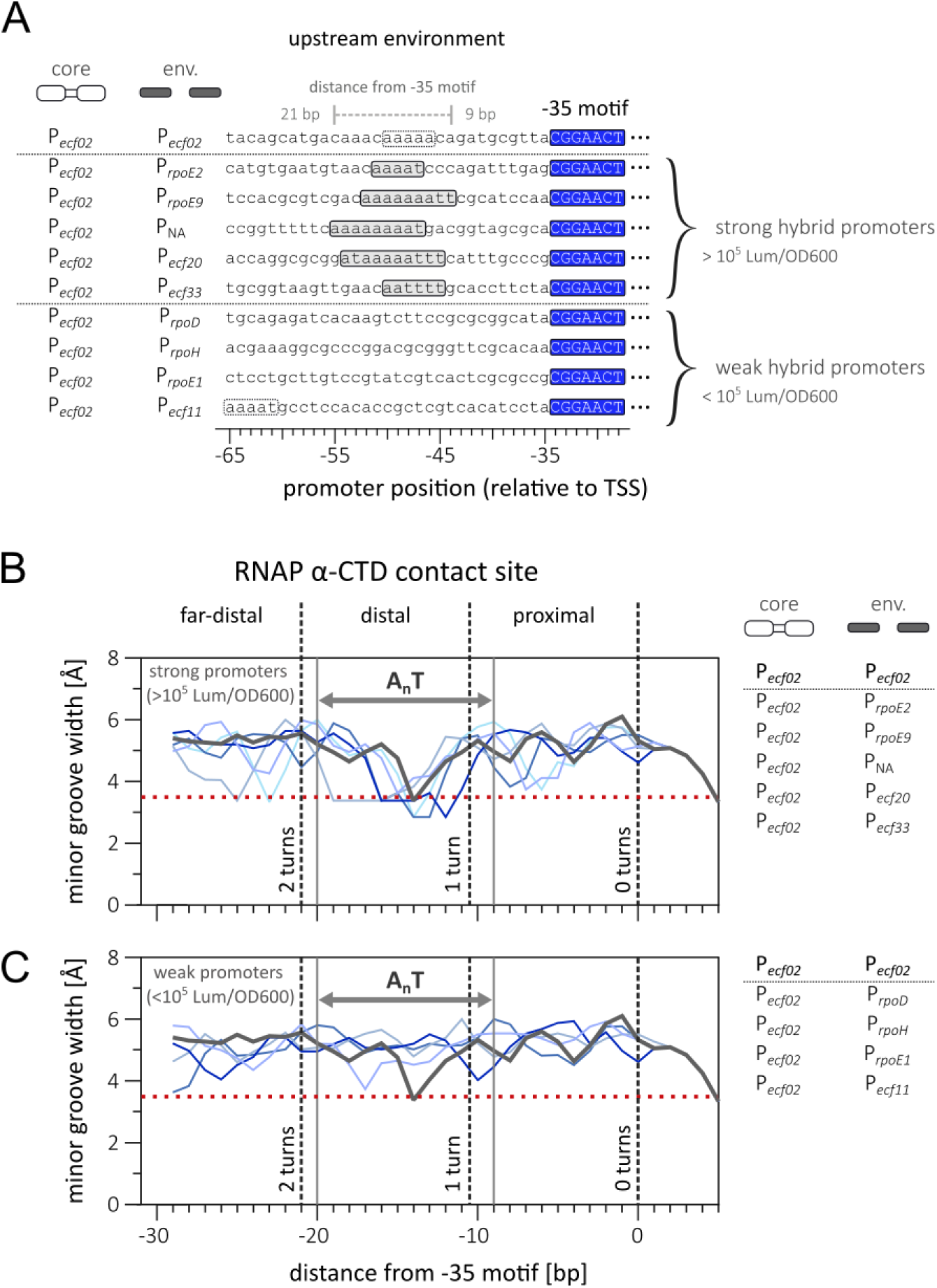
Strong promoter environments contain an UP element-like A_n_T motif. **(A)** Sequences upstream of the P*_ecf02_*-specific −35 box of native P*_ecf02_* and hybrid promoters are shown. In strong promoter environments, an A-rich tract of variable length followed by one or two thymine residues (A_n_T motif; grey boxes) is consistently observed 9 to 20 bp upstream of the 5′ end of the −35 element. This A_n_T motif is positioned approximately one helical turn upstream of the −35 element and is located in a distal UP element position. In one of the weak promoters, an A stretch of similar length without a following T is present in a similar position relative to the −35 element. In another weak promoter, an A_n_T motif is positioned further upstream (about two helical turns upstream of the −35 element). These motifs are indicated by white boxes with dashed outlines. **(B, C)** Predicted minor groove width (MGW) profiles of heterologous sequences (blue lines) upstream of the P*_ecf02_* core promoter for strong (B) and weak (C) promoters. The MGW of the native P*_ecf02_* promoter is shown by dark gray lines in panels. Promoters included in each analysis are listed to the right of the plots. MWG was calculated using DNAshapeR (Chiu *et al*., 2016). Dashed vertical lines indicate successive DNA helical turns upstream of the −35 element and mark the positions of the far-distal, distal, and proximal UP element regions. Solid grey vertical lines denote the location of the A_n_T motif. The horizontal red dashed line marks an MGW of 3.5 Å, the threshold used to predict the presence of an UP element (Todor *et al*., 2020).

To determine whether the presence of an A_n_T motif is predictive of high ECF-dependent promoter activity, we generated a library of hybrid promoters using the weakly active ECF11/P*_ecf11_* switch as a model system. Candidate promoter environments were identified from a dataset comprising 2,729 *S. meliloti* promoters with experimentally defined TSSs (Schlüter *et al*., 2013) (Table S3). Sequence analysis revealed that 17% of these promoters contained an A_n_T motif (Fig. S3A). Ten promoters from this subset were selected at random and used to construct ECF11/hybrid P*_ecf11_* two-plasmid-based switches. ECF switch activities were subsequently quantified in *S. meliloti* (Fig. S3B, C). Seven of the ten hybrid promoters increased the on-state activity above 10^5^ Lum/OD600. Notably, in all seven cases the A_n_T motif overlapped the positions −50 to −48 upstream of the TSS.

Previous work in *E. coli* demonstrated that replacing the promoter sequence upstream of the −35 box with an UP element can increase the activity of heterologous ECF switches containing weak promoters by approximately two orders of magnitude (Rhodius *et al*., 2013). We therefore compared the ability of a commonly used *E. coli* UP element and the A_n_T motif naturally occurring in ECF promoters to enhance the activities of the ECF02/P*_ecf02_* and ECF11/P*_ecf11_* switches (Fig. S4). For this purpose, the upstream promoter environments of P*_ecf02_* or P*_ecf11_* were either replaced by the *E. coli* UP element sequence or modified by introducing the A_n_T motif at positions corresponding to those identified in highly active hybrid promoters. Replacement with the *E. coli* UP element increased the on-state activity of the ECF11/P*_ecf11_*switch by 163%, whereas the same modification reduced the activity of the ECF02/P*_ecf02_* switch by 32% relative to the original promoter. Similarly, implementation of the A_n_T motif increased the activity of the ECF11/P*_ecf11_* switch to levels comparable to those achieved by exchanging both upstream and downstream promoter environments. For example, the A_n_T motif of the P*_rpoE9_* environment alone increased the ECF11-dependent activity of P*_ecf11_* by 158 % whereas the P*_ecf11_* core promoter equipped with both complete P*_rpoE9_*-derived upstream and downstream environments was improved 153 % compared to the native P*_ecf11_*. In contrast, enhancement of ECF02/P*_ecf02_* activity consistently required replacement of both the upstream and downstream promoter environment (Fig. S4, S5).

Taken together, these results suggest that the presence of an A_n_T motif can serve as a reliable predictor of highly active ECF promoter environments when positioned within the distal α-CTD-binding region of the promoter. However, robust enhancement of ECF-dependent promoter activity across different ECF switches required optimization of both the upstream and downstream promoter environments, suggesting that core promoter activity is determined by the combined contribution of upstream and downstream sequence features beyond the A_n_T motif alone.

### Alterations to the A_n_T motif enable tuning of hybrid promoter activities

To further investigate the contribution of the A_n_T motif to promoter activity and assess whether minor sequence changes are sufficient to modulate promoter strength, we designed and characterized targeted mutations of the A_n_T motif in sets of the previously characterized ECF02- and ECF11-dependent hybrid promoters. Two classes of mutations were introduced (Fig. 5). First, TA-swap variants were generated by replacing all adenines with thymines and vice versa within the A_n_T motif. Second, C-disruption variants were constructed by replacing all nucleotides at positions −50 to −48 relative to the TSS − a region overlapping all identified A_n_T motifs − with cytosines.

**Fig. 5:**
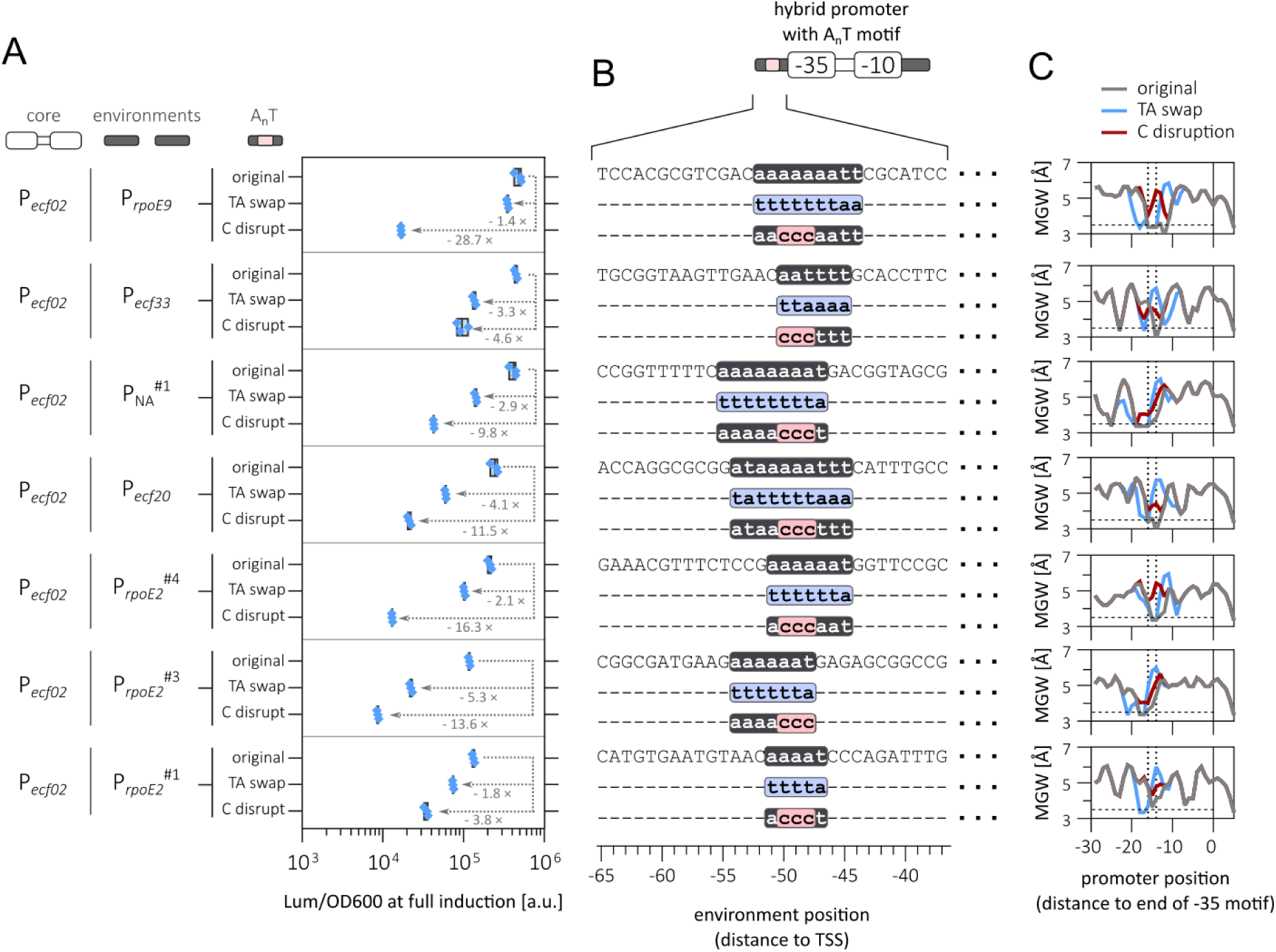
Promoter activities can be modulated through alterations to the UP element-like A_n_T motif. **(A)** Activities of ECF02-dependent hybrid promoters carrying mutations in the UP element-like A_n_T motif, measured in *S. meliloti* Δ*ecf*/Δ*anti-σ* cells 9 h after induction in MOPS minimal medium at full inducer concentration (50 μM). Data represents three biological replicates. Arrows and associated values indicate the fold decrease in promoter activity relative to the corresponding hybrid promoter containing the intact A_n_T motif. **(B)** Upstream sequences of the hybrid promoters showing the native A_n_T motif and its alterations. In the TA-swap mutants, all adenine residues within the A_n_T were replaced with thymine residues and vice versa. C-disruption mutants, positions −50, −49, and −48, which are conserved across all A_n_T motifs examined in this study, were replaced with cytosine residues. Equal bases are designated by dashes “–”. **(C)** Predicted minor groove widths (MGW) profiles of the hybrid promoters and their mutant derivatives, calculated using DNAshapeR (Chiu *et al*., 2016)). Grey lines represent the original hybrid promoters, blue lines the TA-swap variants, and red lines the C-disruption variants. The horizontal dashed lines mark an MGW of 3.5 Å, the threshold associated with the presence of an UP element. The vertical dotted lines indicate the nucleotide positions mutated in the C-disruption variants.

In ECF02-dependent switches, C-disruption mutations of the A_n_T motif reduced promoter activity by 3.8-to 28.7-fold (Fig. 5A). In all cases, these disruptions lowered promoter activity below the threshold of 10^5^ Lum/OD600. TA-swap mutations likewise decreased the activities of ECF02-dependent hybrid promoters, although their effects were generally less pronounced, resulting in activity reductions ranging from 1.4-to 5.3-fold (Fig. 5A). These results suggest that minimal sequence alterations within the A_n_T motif are sufficient to substantially reduce the activity of ECF-dependent promoters.

The reductions in promoter activity were accompanied by changes in the predicted minor groove width (MGW) within the critical region spanning positions −50 to −48. C-disruption variants consistently increased the MGW beyond 3.5 Å (Fig. 5C), whereas TA-swap mutations generally preserved a narrow minor groove. However, TA-swaps occasionally shifted the position of the narrowest groove segments relative to the wild-type sequence. Notably, promoter variants retained high activity only when a narrow MGW was maintained at positions −50 to −48. A similar relationship between promoter activity and MGW was observed for the corresponding ECF11-dependent hybrid promoters (Fig. S6), suggesting that preservation of a narrow minor groove at this location is a general determinant of strong promoter activity.

To further explore the sequence requirements of the A_n_T motif, additional promoter variants were generated in which the motif was repositioned, disrupted by thymine substitutions (analogous to the C-disruption variants), or modified by single-nucleotide substitutions within the critical region at positions −50 to −48. Such mutations were introduced in the P*_ecf02_* and P*_ecf11_* core promoters embedded in the P*_ecf20_* (Fig. S7A) and P*_rpoE9_* (Fig. S7B) environments, respectively. Across both representative promoters, these modifications resulted in a gradual reduction of ECF switch activities by up to 75-fold. Together, these findings demonstrate that the A_n_T motif represents a sensitive and versatile engineering target for the rational tuning of ECF-dependent promoter activity.

### Promoter variant strength is conserved in artificial regulons controlled by an ECF master regulator

To investigate whether promoter variants retain their characteristic activities within a multigene regulatory circuit architecture, we constructed a set of artificial regulons comprising a single ECF master regulator controlling multiple ECF-dependent promoters. Specifically, we used ECF02 as master regulator, and the P*_ecf02_* core promoter with the environments of P*_ecf20_*, P*_rpoE2_*^#3^, and P*_ecf11_*, representing strong, intermediate, and weak promoter variants, respectively (Fig. S9). The regulons consisted of the fluorescent reporter genes *mclover3* and *morange2*, as well as the luminescence reporter cassette *luxCDABE*. We compared two general layouts that shared the input plasmid carrying the salicylate-inducible *ecf02* gene, but differed in the output plasmids. These either carried each a single fusion of one of the ECF02-dependent promoters to one of the reporters (hereafter referred to as 1-regulon configuration) (Fig. 6A) or harbored all three promoter-reporter transcription units (hereafter referred to as a 3-regulon) (Fig. 6B). For the 3-regulon layout, the arrangement and orientation of transcription units, as well as the placement of transcription terminators, were chosen to minimize transcriptional interference and read-through in the output module (Fig. 6B), detailed plasmid maps are shown in Fig. S8). Six distinct 3-regulon output modules were assembled differing in the individual assignment of the three promoter variants to the three reporters (Fig. 6C).

**Fig. 6:**
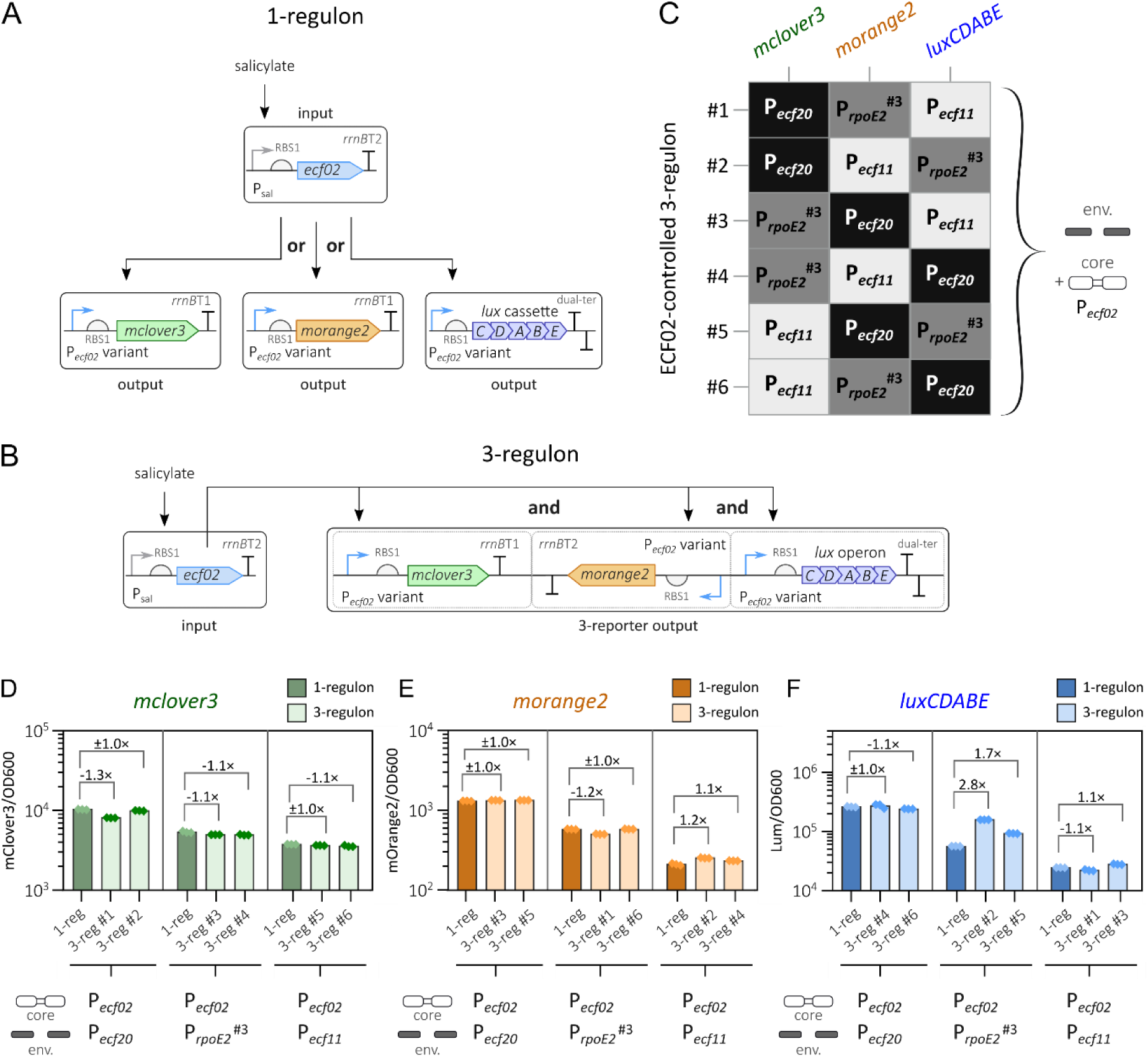
An artificial ECF02-based regulon consisting of three regulated genes. **(A, B)** Schematic of a single-reporter set-up (A) or a 3-regulon (B) using ECF02 as master regulator. The expression of e*cf02* was controlled by salicylate-inducible P_sal_, which was cloned on a single-copy pABCbmodL1 *mob*-derived plasmid. The reporters (*mclover3*, *morange2*, and *luxCDABE*) were paired with one of three different ECF02-dependent promoters of different strength (consisting of the P*_ecf02_* core promoter and environments of P*_ecf20_*, P*_rpoE2_*^#3^, and P*_ecf11_* representing strong, medium, and weak hybrid promoters, respectively). Transcription units containing the reporter genes were cloned on either a single-copy pABCaL1 *mob*-derived plasmid (for the single reporter 1-regulon) or in a head-to-tail-to-head orientation on a single-copy pABCaLM *mob*-derived plasmid (for the 3-regulon). **(C)** Six variants output plasmids of the ECF02-dependent 3-regulon switch were constructed, each containing three distinct ECF02-dependent promoters (no promoter repeated within any single plasmid). Across the collection, each individual promoter was paired exactly two times with the same reporter in different switches. **(D-F)** Reporter signals of *S. meliloti* Δ*ecf*/Δ*anti-σ* strains carrying 1-regulon (dark bars and symbols) or 3-regulon switches (light bars and symbols) captured by the fluorescent reporters *mclover3* (D), *morange2* (E) and the luminescence reporter *luxCDABE* (F) based on three biological replicates. Strains were cultivated in MOPS minimal medium at full induction (50 µM salicylic acid) and relative fluorescence and relative luminescence were measured 23 h and 6 h post induction, respectively. The numbers above the square brackets indicate the average fold difference between promoters in 3-regulon and 1-regulon switches. Positive numbers indicate increased reporter signal measured from strains carrying the 3-regulon compared to those carrying the 1-regulon switch, negative numbers indicate reduced reporter signal measured from strains carrying the 3-regulon switch. A fold change of “± 1.0×” indicates no change in average reporter signal between two switches.

We hypothesized that when compared to the promoter activities in the 1-regulon configuration, each promoter would maintain its characteristic on-state strength in the more complex 3-regulon framework. As a prerequisite for assessing promoter behavior within the regulon context, we first examined whether promoter strengths were affected by the reporter gene used. Across the 1-regulon constructs for all three reporters, the selected promoter variants retained their relative activity ranking Fig. 6D-F, Fig. S10), indicating that promoter-dependent differences were largely reporter-independent.

We next compared promoter activities between the 3-regulon and the corresponding 1-regulon architectures. For nearly all promoter-reporter combinations, on-state expression levels in the 3-regulon setup closely matched those observed in the single-switch configuration (Fig. 6D-F, Fig. S10). Relative reporter activities differed by only −1.3-to 1.2-fold between the two architectures, with the sole exception of the *luxCDABE* reporter controlled by the P_ecf02_ core/P*_rpoE2_*^#3^ environment hybrid promoter. This deviation may reflect context-dependent effects arising from neighboring sequences within the 3-regulon construct.

Analysis of off-state expression revealed no substantial changes for *mclover3* between the 1-regulon and 3-regulon configurations, whereas elevated basal expression was observed for *morange2* and *luxCDABE* in the 3-regulon architecture (Fig. S10). A plausible explanation is cryptic antisense transcription originating from the *morange2–luxCDABE* region of the output plasmid, where no transcription terminators were placed between the oppositely oriented transcription units. These observations highlight the importance of effective transcriptional insulation when designing synthetic multigene regulons.

Collectively, these results demonstrate that ECF-dependent promoters largely preserve their intrinsic transcriptional output when embedded within artificial regulons, supporting the modularity and predictability of this regulatory design strategy.

## Discussion

Alternative σ factors are attractive regulatory components for gene expression control in prokaryotic synthetic biology due to their portability across bacterial hosts. In particular, ECF σ factors, as the simplest members of the σ⁷⁰ family, comprising only the domains required for promoter recognition (Staroń *et al*., 2009; Feklístov *et al*., 2014), have been successfully implemented for orthogonal promoter control across diverse bacterial phyla (Rhodius *et al*., 2013; Pinto *et al*., 2019; Zhao *et al*., 2022; Meier *et al*., 2024). Their high diversity in target promoter specificity (Casas-Pastor *et al*., 2021) and upstream regulatory control, for example via anti-σ factors (Rhodius *et al*., 2013; Boada *et al*., 2025; Wolters *et al*., 2026) provides a powerful resource for scaling synthetic gene network complexity and enabling sophisticated transcriptional programs.

Previous studies have demonstrated the use of alternative σ factors to control genetic circuits (Pinto *et al*., 2018; Boada *et al*., 2025), to tune promoter strength via mutations in core promoter elements (Bervoets *et al*., 2018; Zhao *et al*., 2022), and to control metabolic pathways using σ factor-based master regulators combined with promoter libraries of varying strength (Van Brempt *et al*., 2022). A next step toward constructing synthetic gene networks with a complexity comparable to native bacterial systems is the coordinated control of multiple multi-gene regulons by defined sets of σ factors. Native ECF regulons can comprise more than 80 target promoters (Salgado *et al*., 2024), and many bacterial species encode multiple ECF-dependent regulons (Staroń *et al*., 2009; Cho *et al*., 2014; Lang *et al*., 2018; Casas-Pastor *et al*., 2021; Hatch and Ouellette, 2023), suggesting that the construction of substantially larger synthetic regulatory networks is feasible. However, this approach places stringent demands on promoter-σ factor specificity and requires strict orthogonality to prevent cross-recognition by non-cognate σ factors.

Because promoter specificity is primarily encoded in conserved core promoter motifs (Feklístov *et al*., 2014; Li *et al*., 2019; Lin *et al*., 2019), and mutations in these motifs can alter specificity (Qiu and Helmann, 2001; Domínguez-Cuevas *et al*., 2005), we investigated the upstream and downstream promoter regions as engineering targets to expand the dynamic range of ECF-dependent promoter activity while preserving the core specificity-determining elements. This strategy was motivated by prior observations in *E. coli* showing that inclusion of an UP element can enhance the activity of heterologous ECF-dependent promoters (Rhodius *et al*., 2013). In this study, we show that equipping ECF core promoters with flanking sequences derived from strong donor promoters, either from heterologous sources or the native host, preserves both orthogonality and σ factor specificity of ECF switches. This approach enabled up to 45-fold increase in promoter activity, comparable to improvements achieved through UP element addition (Rhodius *et al*., 2012, 2013) or core promoter mutagenesis (Bervoets *et al*., 2018; Zhao *et al*., 2022). These observations suggest that these strategies for increasing ECF-dependent promoter strength may approach practical upper limits of dynamic range. This limitation is likely imposed by multiple factors, including constraints in core promoter recognition (Rhodius *et al*., 2012, 2013; Li *et al*., 2019; Lin *et al*., 2019), limited availability of RNA polymerase core enzyme (Rhodius *et al*., 2012; Mauri and Klumpp, 2014; Patrick *et al*., 2015), and potential promoter occupancy effects such as RNAP holoenzyme crowding and promoter occlusion (Li *et al*., 2019; Gedeon *et al*., 2021).

In *S. meliloti*, we identified a key contributor to promoter strength in the form of a short UP element-like A_n_T motif upstream of the −35 box. Systematic modification of this motif consistently reduced promoter activity, indicating that it constitutes a distinct and sensitive engineering target for modulating ECF-dependent transcription initiation. In contrast to full-length UP elements that enhance transcription from both RpoD-(Estrem *et al*., 1998, 1999; Ross *et al*., 1998; Presnell *et al*., 2019) and ECF-dependent promoters (Rhodius *et al*., 2013), the A_n_T motif resembles a minimal UP element of approximately 6 bp that can interact with both of the RNAP α-CTDs (Lara-Gonzalez *et al*., 2020). Notably, this motif does not exhibit the typical A/T-rich symmetry characteristic of canonical UP elements (Estrem *et al*., 1998).

Although the A_n_T motif (or canonical UP elements) is important for strong activity of ECF-dependent promoters, it is not sufficient to convert all weak core promoters into strong promoters. One possible explanation is that different ECF-containing RNAP holoenzymes may position the α-CTD differently during transcription initiation, thereby affecting its ability to interact productively with UP element-like sequences (Fig. S11). Consistent with this hypothesis, ECF02- and ECF11-dependent promoters exhibited different sensitivities to positional shifts of the A_n_T motif (Fig. S7), suggesting distinct spatial requirements for α-CTD–DNA interactions. In several cases, increased promoter activity required replacement of the entire promoter context, indicating that additional sequence features also contribute to transcriptional efficiency. These may include DNA structural properties or interactions with downstream promoter elements, such as the CRE, which has been implicated in RNAP interactions during initiation at ECF-dependent promoters (Li *et al*., 2019; Lin *et al*., 2019). A mechanistic understanding of A_n_T-mediated activation and its interplay with other promoter determinants therefore represents an important direction for future work.

Our combined strategy of optimizing the promoter environment through flanking sequence engineering and modulating the A_n_T element enabled the construction of libraries of promoters showing an up to 75-fold difference between activities of the weakest and strongest promoter in *S. meliloti*. Importantly, tuning promoter strength via flanking sequences preserves core specificity determinants while also reducing sequence homology between promoters, thereby potentially lowering the risk of homologous recombination within synthetic regulons.

Importantly, differences in promoter activity observed in the variant libraries were preserved in the context of multi-gene synthetic regulons, supporting the use of ECF switches for rational design of metabolic pathways requiring coordinated expression of genes at different levels. However, increasing regulon complexity will likely exacerbate challenges related to genetic context effects, necessitating attention to design strategies such as transcriptional insulation and careful genomic or plasmid-based positioning of genetic elements.

Overall, ECF switches represent a powerful and increasingly versatile platform for reconstructing the regulatory complexity of bacterial transcriptional networks in synthetic systems. This is supported by their natural diversity, extensive characterization of promoter specificity determinants, and growing availability as standardized tools. Moreover, combining ECF-mediated transcription initiation with additional regulatory layers, including upstream and downstream control mechanisms (Zong *et al*., 2017; Boada *et al*., 2025), enables increasingly sophisticated regulatory logic in such synthetic gene networks. In this context, promoter environment engineering reported in this study provides a transferable, specificity-preserving strategy to expand the dynamic activity range of ECF-dependent promoters and thereby contributes to the scalable design of complex synthetic regulatory systems.

## Methods

### Strains and growth conditions

*E. coli* DH5α (Grant *et al*., 1990) was used for cloning, plasmid maintenance, and as donor for triparental conjugation. Each plasmid listed in Table S4 is associated with an *E. coli* DH5α carrying the respective plasmid. *E. coli* MT616 (Finan *et al*., 1986) was used as conjugation helper strain for triparental conjugation. All *E. coli* strains were grown at 37 °C in LB Lennox (Sambrook *et al*., 1989) (yeast extract 5 g/L, tryptone 10 g/L, NaCl 5 g/L, optional: 1.5 % agar for solid plates). For cultivation in broth, cells were agitated at 200 rpm using an Infors HT (Bottmingen, Switzerland) Ecotron incubator. Selection for and maintenance of plasmids was ensured by adding appropriate antibiotics (gentamicin 10 µg/mL, spectinomycin 100 µg/mL, hygromycin-B 50 µg/mL, or kanamycin 50 µg/mL, chloramphenicol 25 µg/mL, respectively).

All *S. meliloti* strains generated in this study (Table S5) are based on a strain lacking all *ecf* and associated *anti-σ* genes (*S. melilotli* Δ*ecf*/Δ*anti-σ* (Lang *et al*., 2018)). The only remaining σ factors encoded by this strain are the housekeeping σ RpoD, the heatshock σs RpoH1 and RpoH2, and the σ^54^-family σ factor RpoN. *S. meliloti* strains were always cultivated at 30 °C using either complex medium TY (Beringer, 1974) (yeast extract 3 g/L, tryptone 5 g/L, CaCl_2_ · 2 H_2_O 0.4 g/L, optional: 1.5 % agar for solid plates) or a modified 3-(*N*-morpholino)propanesulfonic acid (MOPS) minimal medium (Zhan *et al*., 1991) (basic solution: MOPS 10 g/L, mannitol 10 g/L, sodium glutamate 3.55 g/L, MgSO_4_ · 7 H_2_O 0.246 g/L, adjusted to pH 7.2 using KOH). After sterilization, the basic solution was supplemented with each 1 mL/L of CaCl_2_ (250 mM), FeCl_3_ · 6 H_2_O (10 g/L), biotin (1 mg/mL in 0.1 M NaOH), oligo element solution (composition below), and 2 mL/L K_2_HPO_4_ (1 M). Oligo element solution consisted of H_3_BO_3_ (15 g/L), MnSO_4_ · 4 H_2_O (11.15 g/L), ZuSO_4_ · 7 H_2_O (1.435 g/L), CuSO_4_ · 5 H_2_O (0.625 g/L), CoCl_2_ · 6 H_2_O (0.325 g/L), and NaMoO_4_ · 2 H_2_O (0.6 g/L). *S. meliloti* cultured in broth were grown either in glass tubes, transparent 96-well multi well plates (MWPs) or black MWPs. When grown in glass tubes, *S. meliloti* cultures were aerated at 200 rpm using Infors HT Multitron shaker. Cultures grown in transparent MWPs (Greiner Bio-One, Kremsmünster, Austria; catalog number: 655161) were kept to a maximum volume of 200 µL, sealed with a non-breathable transparent foil (Greiner Bio-One, catalog number: 676001), and agitated at 1200 rpm using an Heidolph (Schwabach, Germany) Titramax 1000 incubator. Cultures were grown in black MWPs (Greiner Bio-One, catalog number: 655096) were used for plate reader experiments and cultivation is described in the respective chapter. *S. meliloti* strains used in this study are naturally resistant to streptomycin at a concentration of 600 µg/mL. When appropriate, further antibiotics were used for plasmid selection and maintenance (gentamicin 30 µg/mL for solid plates, 20 µg/mL for broth; spectinomycin 200 µg/mL; hygromycin-B 100 µg/mL; or kanamycin 200 µg/mL, respectively).

### Design of synthetic hybrid promoters

Alternative ECF02- and ECF11-dependent hybrid promoters were created by combining the core promoter sequences (−35 box/spacer/-10 box) from base promoters P*_ecf02_* and P*_ecf11_*, respectively, and upstream and downstream environmental sequences of donor promoters. Donor promoter sequences originated from *S. meliloti*, *E. coli*, *Pseudomonas fluorescens*, *Pseudomonas synringae*, and *Bradyrhizobium japonicum* (Table S2). When the −35 and −10 boxes of the donor promoter were known, care was taken to completely remove all sequences belonging to these boxes from the resulting hybrid promoter by including more nucleotides from the base promoter in the final design. A detailed description of the process is depicted in Fig. S2.

### Manipulation and extraction of DNA

Plasmids used throughout this study were constructed using restriction/ligation-based and Golden Gate cloning methods (Sambrook *et al*., 1989; Weber *et al*., 2011; Werner *et al*., 2012). Standard restrictions were performed with 300 to 600 ng plasmid DNA, digested with Thermo Scientific™ (Waltham, Massachusetts, USA) FastDigest restriction enyzmes (catalog numbers FD0684 and FD1564) and purified using Omega Bio-Tek (Norcross, Georgia, USA) E.Z.N.A.® Cycle Pure Kit (catalog number D6492) following manufacturer’s instructions. For the annealing and phosphorylation of oligo nucleotide pairs, 100 fmol per oligo nucleotide (synthesized by Integrated DNA Technologies, Inc. (IDT), Coralville, Iowa, USA) (Table S6) were phosphorylated using 5 U of T4 polynucleotide (Thermo Scientific™, catalog number EK0031) and applied in a final concentration of 10 fM for cloning reactions. Ligations were performed with 5 U T4 DNA ligase (Thermo Scientific™ catalog number EL0014). Gene fragments for the coding sequences of *mclover3* and *morange2* were ordered as double stranded gene fragments (Twist Bioscience, San Francisco, California, USA) and are listed in Table S6. Golden Gate reactions were performed with 15 fmol per DNA part and 5 U BsaI (Thermo Scientific™, catalog number ER0292) or BpiI (Thermo Scientific™, catalog number ER1012) and 2.5 U T4 DNA ligase (Thermo Scientific™, catalog number EL0014) in T4 DNA Ligase Buffer (catalog number B69). Each Golde Gate reaction was incubated at 37 °C for at least five hours before heat inactivation (10 min at 50 °C followed by 10 min at 80 °C). Plasmids were isolated from overnight culture of *E. coli* DH5α grown in LB broth using Omega Bio-tek E.Z.N.A.® Plasmid DNA Mini Kit (catalog number D6943) according to manufacturer’s instructions. Plasmids were verified via PCR with Taq polymerase (NEB, Ipswich, Massachusetts, USA; catalog number M0273) or Sanger sequencing (Microsynth Seqlab GmbH, Göttingen, Germany). Detailed descriptions of cloning strategies for all plasmids are given in Table S4.

### DNA transfer to *E. coli* and *S. meliloti*

Chemically competent *E. coli* DHα were prepared using the Inoue method (Inoue *et al*., 1990) and transformed following a standard heat shock protocol (Sambrook *et al*., 1989). Colonies were visible after 16 to 18 hours incubation at 37 °C and single colonies were used for validation of assembled plasmids.

To simultaneously introduce up to three mobilizable *repABC*-based plasmids into *S. meliloti*, we performed triparental conjugation (Datta *et al*., 1971) using *E. coli* MT616 conjugational helper strain (Finan *et al*., 1986) and *E. coli* DH5α carrying respective plasmids as parental strains. Briefly, *S. meliloti* recipient strains, *E. coli* MT616, and up to three *E. coli* DH5α donor strains were grown overnight on respective agar plates. All strains were mixed in approximately equal ratios in LB or TY without antibiotics. 5 to 20 µL of mixed suspensions were spotted onto TY agar plates without antibiotics and incubated at 30 °C overnight. Mating spots were resuspended in 500 µL sterile 0.9 % NaCl and plated in serial dilutions onto solid TY plates containing streptomycin and respective antibiotics for selection of plasmids. Visible colonies formed after three days of incubation at 30 °C.

### Long-term storage of bacterial cultures

For long-term storage of *E. coli* strains, respective strains were streaked on one half agar plate with appropriate antibiotics and resuspended in 1 mL LB with 15 % glycerol without antibiotics. *S. meliloti* cultures intended for long term storage were grown in 96-well MWPs in TY broth with antibiotics. DMSO was added to a final concentration of 75 µg/mL. Long-term cultures were stored at −80 °C.

### Plate reader assays

Plate reader assays with *S. meliloti* shown in all figures except Fig. 6 and Fig. S10 were performed using a laboratory automation facility described elsewhere (Meier *et al*., 2024). In brief, cells were grown in transparent MWPs in TY broth with antibiotic selection. Overnight cultures were diluted 1:500 in MOPS minimal medium and growth was allowed for approximately 18 hours. OD600 was synchronized using a Tecan (Männedorf, Switzerland) Freedom Evo200 liquid handler platform and cultures were transferred to black MWPs with removable lids with condensation rings (Greiner Bio-One, catalog number: 656170). Inducer was added after an initial measurement. Plates were agitated every hour on a Thermo Scientific™ Teleshake with 2 mm amplitude and OD600 and luminescence (integration time: 250 ms) were measured using a Tecan Infinite M200 reader integrated in the liquid handling platform every 3 h for a duration of 12 h total. Data shown in Fig. 6 and Fig. S10 were generated as follows: *S. meliloti* strains were inoculated in TY broth with antibiotics in glass tubes and grown overnight. After dilution to OD 0.05, strains were distributed to black MWPs and salicylate was added to a final concentration of 0 µM and 50 µM, respectively. OD600 and luminescence (integration time: 250 ms) was captured after six hours of growth using a Tecan Infinite M200 Pro reader. OD600 as well as mClover3 fluorescence (excitation 485 nm, emission: 520 nm, gain: 82) and mOrange2 fluorescence (excitation: 530 nm, emission: 565 nm, gain: 82) was captured after twenty hours using the same machine. Relative fluorescence and relative luminescence were calculated as such: Raw OD600 and raw fluorescence or luminescence values were corrected by blank value subtraction. When blank-corrected fluorescence or luminescence reached values below 0, these were set to 0 instead. Relative fluorescence and relative luminescence were obtained by normalizing blank-corrected fluorescence or luminescence to blank-corrected OD600.

### Statistical methods

Quantitative data mentioned throughout the manuscript represents average values of at least three biological replicates. Biological replicates were identified as individual colonies from the same conjugation plate. Errors are represented as ± S.D. Statistical significance was calculated as multiple *t* test using the Holm-Sidak method (GraphPad Prism v8.0.1). Dose-response curves of inducible promoters were fitted with GraphPad Prism v8.0.1 using the non-linear fit function (variable slope, four parameters).

## Author Contribution

A.B. and D.M. conceived the study. C.R. and D.M. designed the experiments. C.R., V.S., and D.M. performed the experiments. C.R. and V.S. analyzed the data. E.M.E. modelled protein complex/DNA interactions and presented the results of the model in Fig. S11. C.R. created the majority of figures. A.B. and C.R. wrote the manuscript, and all authors revised and approved the manuscript.

## Declaration of competing interests

The authors declare no competing interest.

## Reporting summary

Further information on research design is available in the Nature Portfolio Reporting Summary linked to this article.

## Data availability

The numerical source data for the graphs can be found in Supplementary Data 1 and 2.

## Supporting information

Supplemental Data 1, numerical values for the generation of figures in the main manuscript

Supplemental Data 2, numerical values for the generation of supplemental figures

## Acknowledgements

We thank Patrick Manz and Dr. Javier Serrania for assistance with the SYNMIKRO laboratory automation core facility used for plate reader experiments.

## Funding statements

A.B. discloses support for the research of this work from the German Federal Ministry of Education and Research (grant 031L0010B) and the German Research Foundation (EXC 3048/1, project no. 533620160).

## Supplementary information

Supplemental information is provided as additional files.

## Notes

### Competing Interest Statement

The authors have declared no competing interest.

