## Supplemental Table S1, Supplemental Figures S1 to S11 for "Dynamic expression range expansion of ECF sigma factor-dependent synthetic regulons by promoter context engineering"

*Running title:* Promoter environments enable artificial regulons

Christian Rauch<sup>a, b, c</sup>, Doreen Meier<sup>a, b</sup>, Vinca Seiler<sup>b</sup>, Eslam M. Elsayed<sup>a, b</sup>, Anke Becker<sup>a, b, c\*</sup>

<sup>a</sup> Center for Synthetic Microbiology (SYNMIKRO), Philipps-Universität Marburg, Karl-von-Frisch-Straße 14, 35032 Marburg, Germany

<sup>b</sup> Department of Biology, Philipps-Universität Marburg, Germany, Karl-von-Frisch-Straße 8, 35032 Marburg, Germany

<sup>c</sup> Microbes-for-Climate (M4C) Cluster of Excellence, Philipps-Universität Marburg, Karl-von-Frisch-Straße 14, 35032 Marburg, Germany

Separate files:

**Table S2** Sequences of ECF-dependent promoters

**Table S3** Hybrid promoter library based on Schlüter mTSS

**Table S4** List of plasmids and associated *E. coli* DH5α strains

**Table S5** List of *S. meliloti* strains

**Table S6** List of oligo nucleotides and synthetic dsDNA fragments

**Data S1** Data used to generate main figures

**Data S2** Data used to generate supplementary figures

23 **Supplementary Tables**

24 **Table S1:** Protein sequences of ECFs used in this study. Given are names used throughout this manuscript and the
25 phylogenetic groups and subgroups associated with the ECF according to current nomenclature (Casas-Pastor *et*
26 *al.*, 2021).

| Name used<br>in this study | Phylogenetic<br>group | Sequence |
| --- | --- | --- |
| ECF02 | ECF02 s1 | MSEQLTDQVLVERVQKGDQKAFNLLVVRYQHKVASLVSRYVPSGDVPDVV<br>QEAFIKAYRALDSFRGDSAFYTWLYRIAVNTAKNYLVAQGRRPPSSDVDA<br>IEAENFESGGALKEISNPENLMLSEELRQIVFRTIESLPEDLRMAITLRE<br>LDGLSYEEIAAIMDCPVGTVRSRIFRAREAIDNKVQPLIRR |
| ECF11 | ECF11 s5 | MMSDSPQKLGRNEWNAYMDKVKAKDREAFAFVFRFYAPKLKQFAYKHVGN<br>EQVAMEMVQETMATVWQKAHLYDGKKSALSTWIYTIIRNLCFDLLRKQKG<br>KELHIHSDDIWPSEYYPPDMVDHYSPEQDMLKEQVVKFLDILPKNQRDVL<br>QAVYLEELPHQQVAELFDIPLGTVKSRLRLAVEKLRHSMHTEQL |
| ECF20 | ECF290 s1 | MNETDPDLELLKRIGNNDAQAVKEMVTRKLPRLLALASRLLGDADEARDI<br>AQESFLRIWKQAASWRSEQARFDTWLHRVALNLCYDRLRRRKEHVPVDSE<br>HACEALDTRPAPDEQLEASAQSRRMAQALDQLPDRQREAIVLQYYQELSN<br>TEAAALMQISVEALESLLSRARRNLRSHLAEAPGADLSGRRKP |
| ECF33 | ECF33 s1 | MSTKQAATDDVLIARIAQGDRVAMQVLYGRHHVKVFRFGLRLVRNEQIAE<br>DLISEVFLDVWRQAGKFEGRSSVSTWLLAITRFKALSALRRRKDAELDDE<br>AAAAIEDPSDDPEIAVQKKDTGEALRKCLSSLTPEHREIVDLVYYHEKSI<br>EEVAEIVGIPENTVKTRLFYARKKLADLLQAAGVQRGWP |
| ECF31 | ECF31 s1 | MDTQEEQRLIQQAKEGNDEAFTALFHYHYSFLYKYLLKLSLHPDLSEELV<br>QETFLKGYIHLRSFQGRSKFSTWLISIASRLYLDHQKKRKREWKRNQTVT<br>EETIRKIKWDVSAKGAEWSETLDLFSKLDPKLRTPVLLRHYYGYTYAEIG<br>VMLQIKEGTVKSRVHKGLQQIRKEWDDE |
| ECF15 | ECF15 s1 | MTQTPKAPAKHRDPRDELPEHLPALRAFAISLTRNVAVADDLVQDTIVKA<br>WTNFDKFTEGTNLRAWLFTILRNTFYSDKRKHRREVPDPEGVHAASLFVK<br>PAHDGHLAFSDFSAAFDQLSPEHREVLILVGASGFAYEDAAQMMGVAVGT<br>VKSRANRARARLAELLGLEKGEEIFSGVDGQTLAVMSRSGMTAA |
| ECF42 | ECF41 s1 | MAATDISALLDTLWRREASRIIGALARQLRDVGLAEELAQDALVAALEHW<br>PRNGIPDNPAAWLMTTAKHRAIDRLRHYQLQRRKQDALSFEIEREQQAAS<br>AAQLVDPDDDLGDDQLRLMFVACHPALGTDARVALTLRLLGGLSTDAIAR<br>AFLVPEATIAQRIVRAKRTLSDKQVPFEVPRGSERHARLASVLEVLYLIF<br>NEGYAASDGDDAQRPALCHEALSLIGVLAEQMPTAAEVHALRALMALQAS<br>RSAARSDASGAPVLLLEQDRSRWHQPLIQLGLAALARAQQLGGGESSYAL<br>QAAIAACHVTAAQAQDTDWPRIAALYTRLAQRVPSPVIALNRAVAIAMAE<br>GPAAGLVLVEALQQDSALRHYHLLPSVRGDLLYKLGRFDEARADFLHAAT<br>LAGNARDKTFLLTRADACSGLPA |

**Supplementary information: Promoter environments enable artificial regulons**

|  |  |  |
| --- | --- | --- |
| ECF26 | ECF26 s5 | MNDLDPVVTVSDGICALLPRLRRFARAIAGHPADADDLLQVAIERALRHC<br>AQWRPETPLQYWLFGIIRHAWLDEVRAQRRRRQLFVSAAEGEQVGDS PME<br>RQQEWMAVQAAMAQLPDEQRWPIALVLIEGLSYRDAAAVLEIPIGTLTSR<br>LARGREALQALLEEKs |
| ECF34 | ECF19 s4 | MNTRTRPTTTPVPPVEETRYEEELAHGLVKADEDAFAAIYRRWGSLVHTL<br>ATRS LGDAHEAEDVTQQVFVGAWRGRHGFRPERGTLGAWLVGITRRKVVD<br>ALAARTRRLSLVESAGQDITPARLVQPALDEV LDRVLLVEALSRLPQAQR<br>DVLCMAFYEDLTQAQIAERTGVPLGTVKSHARRGLHRLRTAVGPGAAHDT<br>CV |
| ECF17 | ECF17 s4 | MARVSGAAAAEAALMRALYDEHAAVLWRYALRLTGDAAQAEDVVQETLLR<br>AWQHPEVIGDTARPARAWLFTVARNMIIDERRSARFRNVVGSTDQSGTPE<br>QSTPDEVNAALDRLLIADALAQLSAEHRAVIQRSYYRGWSTAQIATDLGI<br>AEGTVKSRLHYAVRALRLTLQELGVTR |

28 **Supplementary Figures**

29

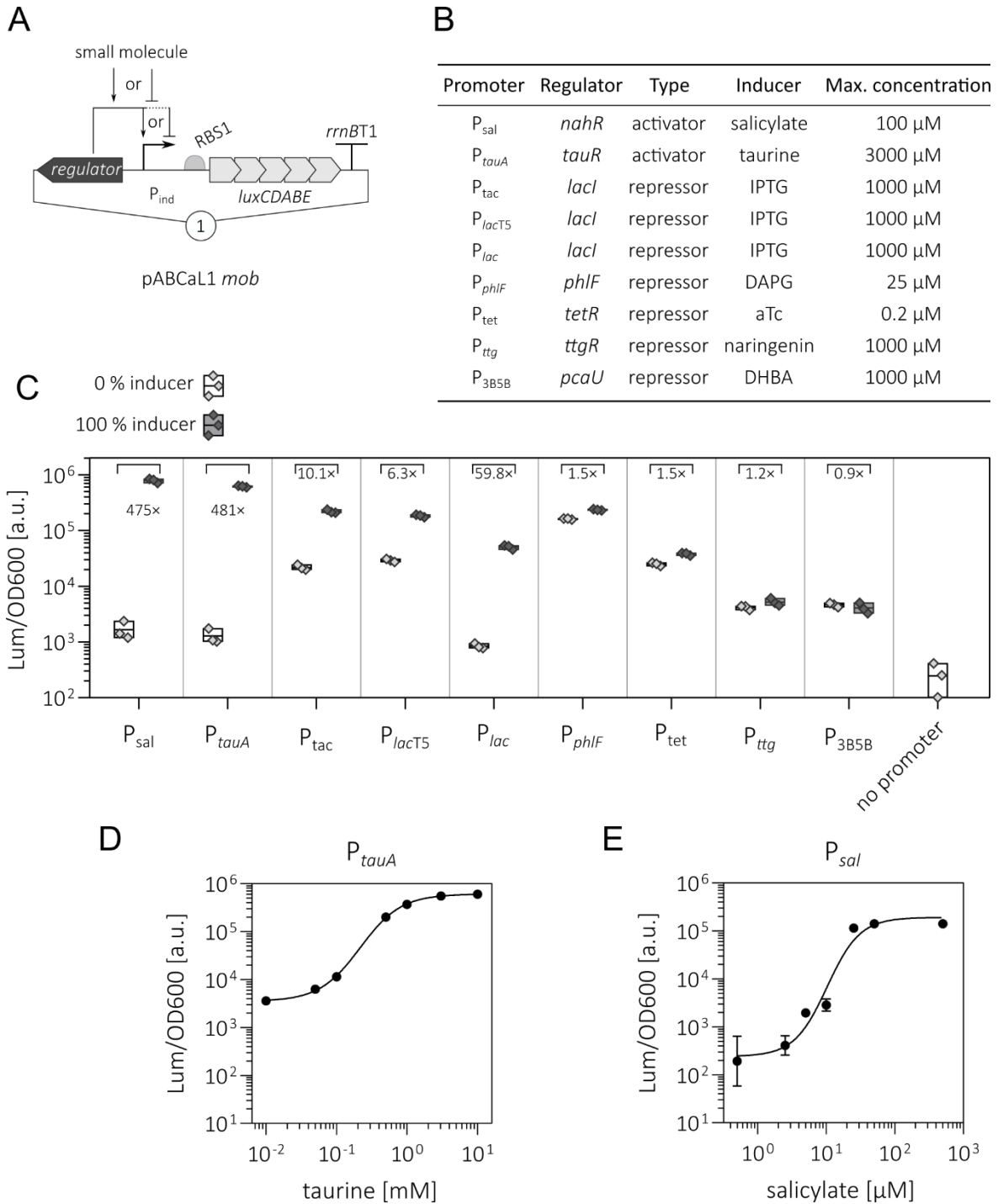

**Figure S1: Characterization of novel inducible promoters in *S. meliloti* Δecf/Δanti-σ.**

(A) Schematic representation of report plasmids for the characterization of inducible promoters. A single copy plasmid pABCaL1mob carried promoter/regulator pairs which controlled the transcription of the *luxCDABE* reporter cassette with a standardized 5'-UTR upstream of *luxC*. Notably, the taurine-inducible system $P_{tauA}$ relied on the TauR regulator protein endogenously encoded by *SMb21525* on an endogenous chromid. Regulators comprised a set of transcriptional repressors and activators. (B) Inducible promoters were induced with sodium salicylate, taurine, isopropyl-β-D-thiogalactopyranosid (IPTG), 2,4-diacetylphloroglucinol (DAPG), anhydrotetracycline (aTc), naringenin, or dihydroxybenzoic acid (DHBA) with the maximal inducer concentrations given.

**(C)** Luminescence output of promoter/regulator reporter plasmids in *S. meliloti* $\Delta ecf/\Delta anti-\sigma$ in MOPS-buffered minimal medium 9 h after the addition of respective inducers. The “no promoter” strain carried an empty pABCaL1mob plasmid. The maximal inducer concentrations (100 %) differed per substance and promoter/regulator pair and is given in (B). The numbers indicate the fold change between the on-state (presence of 100 % inducer) and the off-state (0 % inducer). **(D, E)** Dose response of the $P_{touA}$ reporter plasmid (D) or the $P_{sal}$ reporter plasmid (E) in *S. meliloti* $\Delta ecf/\Delta anti-\sigma$ cultured analogous to (C). Error bars represent one standard deviation. Sigmoidal relationship between luminescence output and inducer concentration was found by fitting the data using GraphPad Prism v8.0.1 (non-linear fit, variable slope (four parameters)). Each strain in (C-E) was measured in at least three biological replicates.

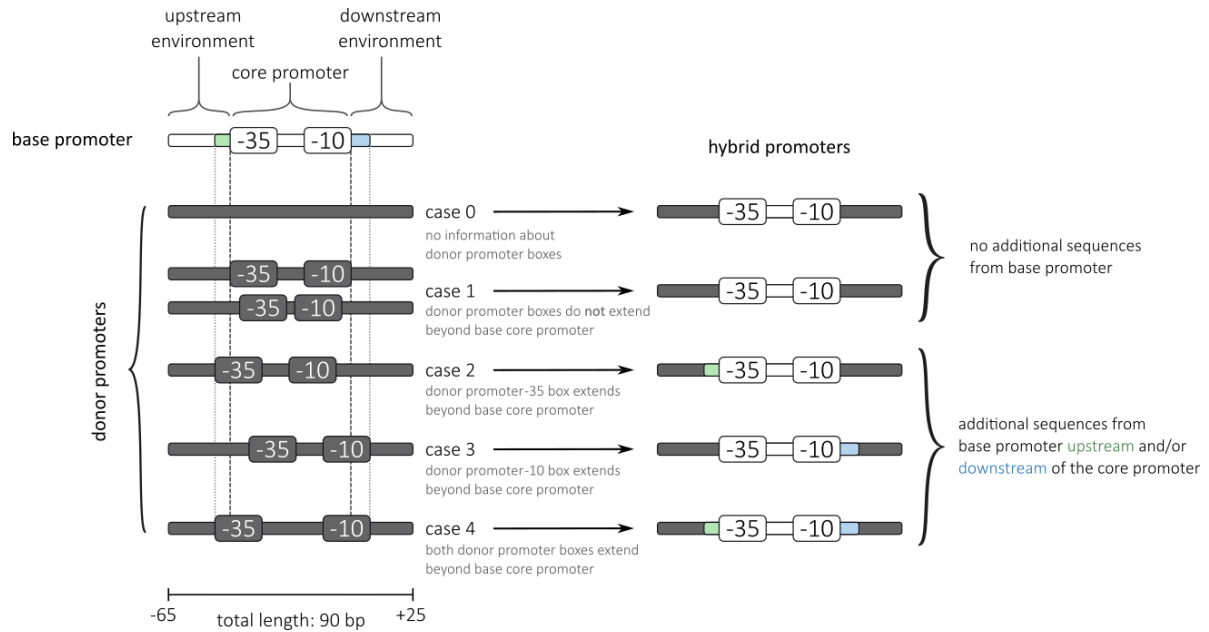

**Figure S2: Design of hybrid promoters.**

Promoters used in this study ranged from position -65 to +25 relative to the TSS. We extended the previously used size of promoter fragments (-60/+20) (Rhodius *et al.*, 2013; Meier *et al.*, 2024) for the following reasons: In *S. meliloti*, mRNAs with long 5' UTRs are common with their distribution peaking at around 25 nt downstream of the TSS, yet extending well beyond to up to 309 nt (Schlüter *et al.*, 2013). We therefore included more base pairs downstream of the TSS. Since the -35 and -10 core promoter boxes are spaced differently for different $\sigma$ factor classes, putative elements in the upstream region may also be preserved in a slightly longer promoter fragment (Rhodius *et al.*, 2012).

Each promoter sequence (90 bp, spanning -65 to +25 relative to the transcription start site) was segmented into the core promoter (-35 box, spacer, and -10 box) and flanking upstream and downstream environments. Hybrid promoters were constructed by combining the core promoter from a base promoter with upstream and downstream environments of a donor promoter. All donor-derived core promoter sequences were systematically removed from the hybrid constructs whereby the positioning of base and donor core promoters relative to each other determined whether additional base promoter-derived sequences needed to be retained. We classified hybrid constructs into five cases: (0) donor core promoter motifs undefined, no additional base sequences retained; (1) donor and base core promoters aligned or base core promoter extends beyond donor core promoter, no additional base sequences retained; and (2-4) donor -35 and/or -10 boxes extend beyond base core promoter boundaries, requiring retention of additional base-derived flanking sequences to eliminate all residual donor core promoter nucleotides. Core promoter boundaries are indicated by black dashed lines (first base of -35 to last base of -10 box), with grey dotted lines representing donor core promoter extent when exceeding base coordinates. Retained base-derived sequences are highlighted in green and blue.

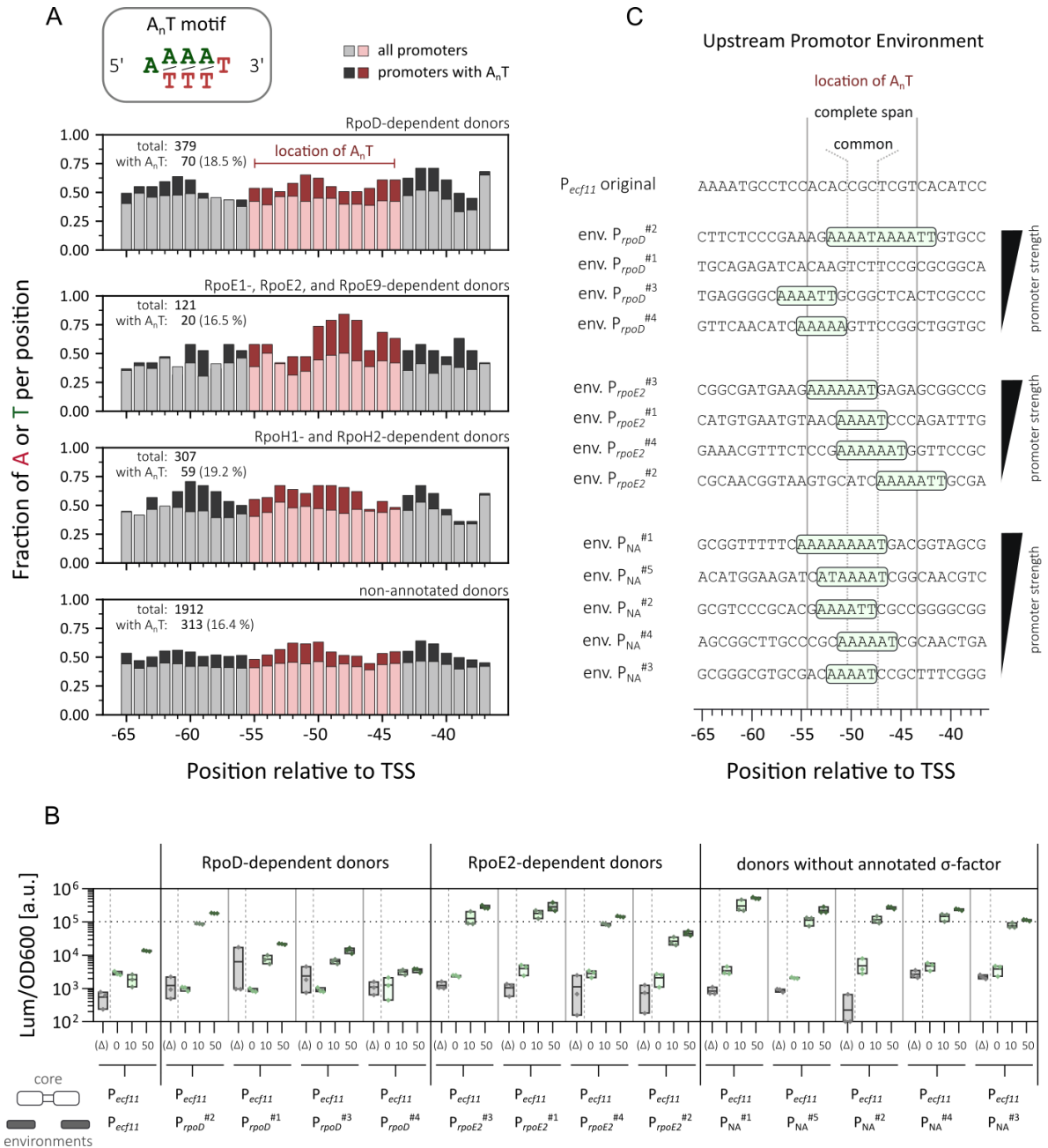

**Figure S3: A library of hybrid ECF-dependent promoters based on transcription start site data.**

**(A)** Analysis of the presence of an A<sub>n</sub>T motif in a library of ECF11-dependent environmental hybrid promoter based on endogenous *S. meliloti* promoters from a set of transcription start sites strongly associated with mRNAs (Schlüter *et al.*, 2013). The library contained a total of 2729 hybrid promoters and can be found in Table S3. Upstream environments that contained an adenine and thymine stretch of at least five consecutive bases starting with adenine and ending with thymine were designated as containing an A<sub>n</sub>T motif. General distribution of adenine and thymine presence at defined distances from the TSS is shown for all promoters (light bars) or those containing an A<sub>n</sub>T motif (dark bars). Red bars indicate the location of experimentally verified A<sub>n</sub>T motifs (positions -55 to -44, Fig. 4). **(B)** Luminescence output of ECF11/*P<sub>ecf11</sub>*-based two-plasmid switches in *S. meliloti* Δecf/Δanti-σ in MOPS-buffered minimal medium 9 h after the addition of 0 μM, 10 μM, or 50 μM salicylic acid. Strains lacking the *ecf11* gene on the input module plasmid are designated as “($\Delta$)”. ECF11-dependent hybrid promoters were chosen at random based on the presence of an A<sub>n</sub>T motif in their upstream region. For comparison, previously characterized ECF11/*P<sub>ecf11</sub>* switches (Fig. 2C) are also shown (environments labelled with “#1”) and may therefore lack an A<sub>n</sub>T motif. Black solid vertical lines separate hybrid promoters by $\sigma$ factor annotation of the donor environments. Grey solid vertical lines separate each individual switch. Grey dashed vertical lines separate “no *ecf*” control strains from strains with an active switch. Switches with an on-state activity above 10<sup>5</sup> Lum/OD600 (indicated by the dashed horizontal line) were considered strong promoters. **(C)** Sequences of the upstream environments of promoters from switches characterized in (B). The A<sub>n</sub>T motif of each switch is highlighted with a green box. The location of all A<sub>n</sub>T motifs experimentally verified previously is given by the solid grey lines. Three positions common to all A<sub>n</sub>T motifs (-50

to -48 bp relative to the TSS) are indicated by dotted lines. Notably, only promoters whose A<sub>n</sub>T motif overlapped the common three positions were characterized as strong promoters.

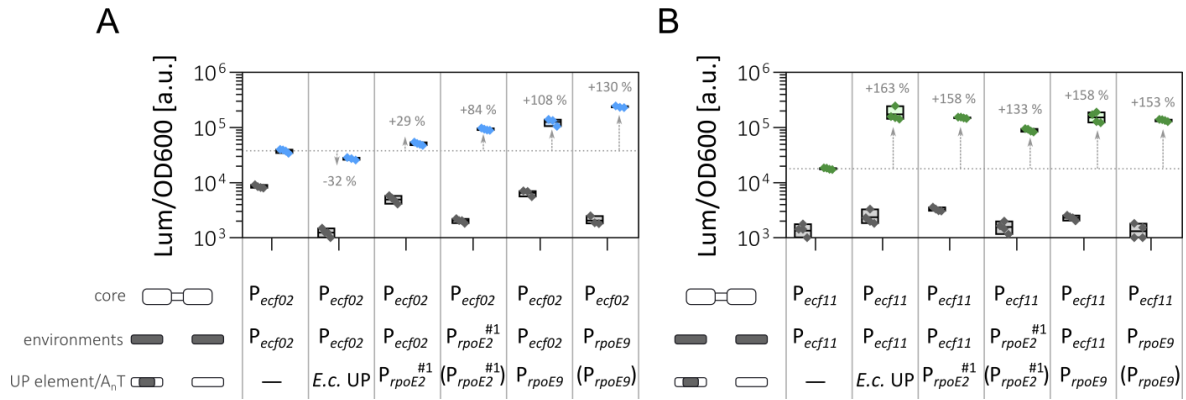

**Figure S4: Functionality of an *E. coli* UP element in heterologous ECF-dependent promoters *S. meliloti*.**

Characterization of ECF02/P<sub>ecf02</sub>- (A) and ECF11/P<sub>ecf11</sub>-based (B) two-plasmid switches in *S. meliloti* $\Delta ecf/\Delta anti-\sigma$ in MOPS-buffered minimal medium 9 h after the addition of 0 μM (off-state, grey boxes and diamonds) or 50 μM (on-state, blue or green boxes and diamonds) salicylate. ECF-dependent promoters were either the native promoter, had an added A<sub>n</sub>T motif or UP element as used by Rhodius *et al.* (2013), or were complete upstream and downstream environmental hybrid promoters. At least three biological replicates per strain are depicted. The dashed horizontal line shows the average on-state activity of the native ECF02-dependent (A) or ECF11-dependent (B) promoter. Numbers and arrows indicate the relative change of on-state activity of modified promoters compared to the respective native promoters.

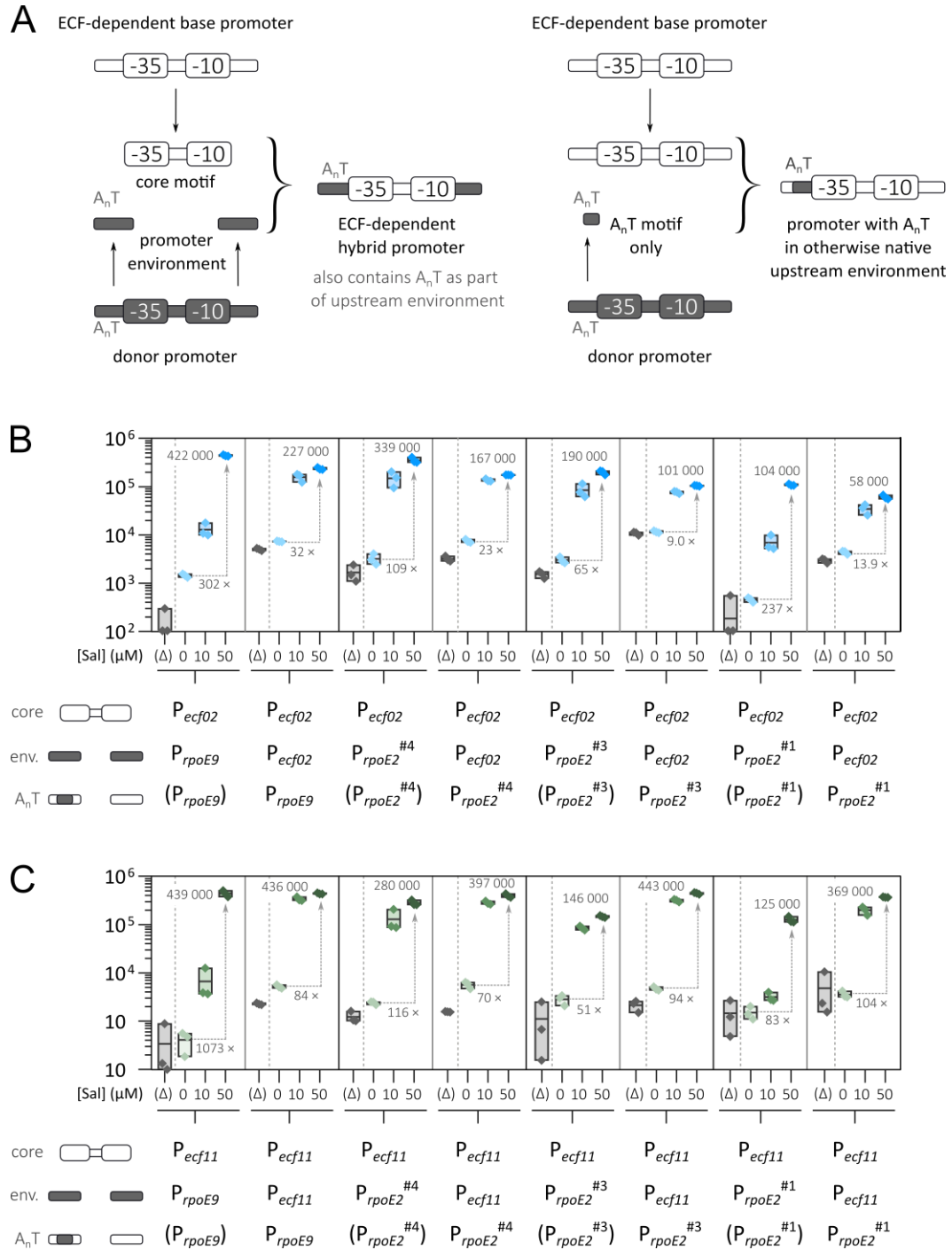

**Figure S5: Functional characterization of the A<sub>n</sub>T motif as a module in ECF-dependent promoter engineering.**

**(A)** Schematic representation procedure employed to engineer hybrid ECF-dependent promoters. Next to previously established environmental hybrid promoters that carry the A<sub>n</sub>T motif as part of their upstream environments, the A<sub>n</sub>T motif itself was used to replace respective bases in the upstream environment of a base promoter. Other environmental sequences and the core promoter were native. Respective promoter sequences are listed in Table S2. **(B, C)** Luminescence output of ECF02 - **(B)** and ECF11-dependent **(C)** two plasmid-based switches in *S. meliloti* Δ*ecf/anti-σ* in MOPS-buffered minimal medium 9 h after the addition of 0 μM, 10 μM, or 50 μM salicylate. The corresponding “no *ecf*” control strain for each switch is denoted by “(Δ)”. Solid vertical lines separate switches and the dashed vertical line separates complete switches from “no *ecf*” control strains. The output module of the switches contained promoters that were either complete upstream and downstream environmental hybrids or only carried the A<sub>n</sub>T motif of the respective donor promoter in their upstream environment. The numbers on the grey arrows denote the average fold change between the off- and the on-state of each switch. Numbers at the boxes for the cultures induced with 50 μM salicylate indicate average on-state activity of the respective switch.

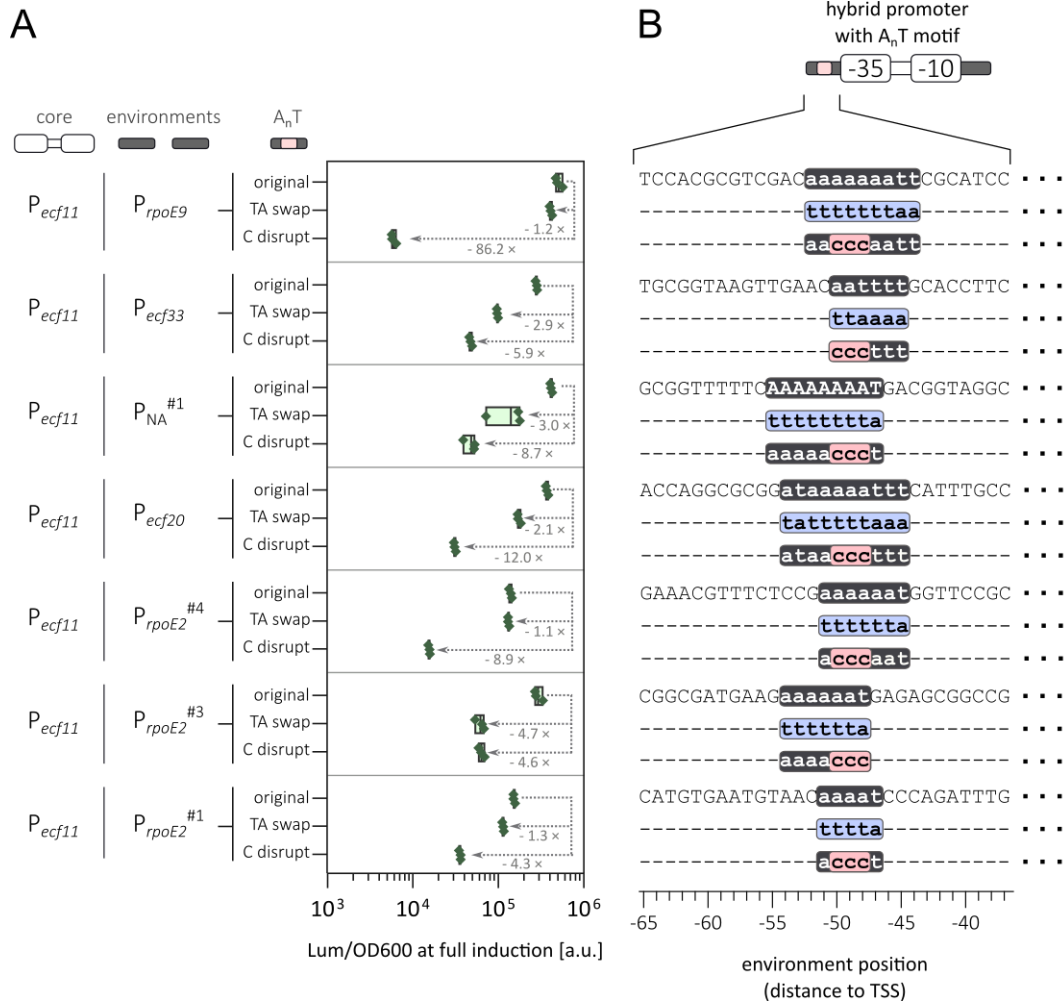

**Figure S6 Promoter activities can be modulated through alterations to the UP element-like A<sub>n</sub>T motif.**

**(A)** Activities of ECF11-dependent hybrid promoters carrying mutations in the UP element-like A<sub>n</sub>T motif, measured in *S. meliloti* $\Delta ecf/\Delta anti-\sigma$ cells 9 h after induction in MOPS-buffered minimal medium at full inducer concentration (50 μM). Data represents 3 biological replicates. Arrows and associated values indicate the fold decrease in promoter activity relative to the corresponding hybrid promoter containing the intact A<sub>n</sub>T motif. **(B)** Upstream sequences of the hybrid promoters showing the native A<sub>n</sub>T motif and its alterations. In the TA-swap mutants, all adenine residues within the A<sub>n</sub>T were replaced with thymine residues and vice versa. C-disruption mutants, positions -50, -49, and -48, which are conserved across all A<sub>n</sub>T motifs examined in this study, were replaced with cytosine residues. Equal bases are designated by dashes “-”.

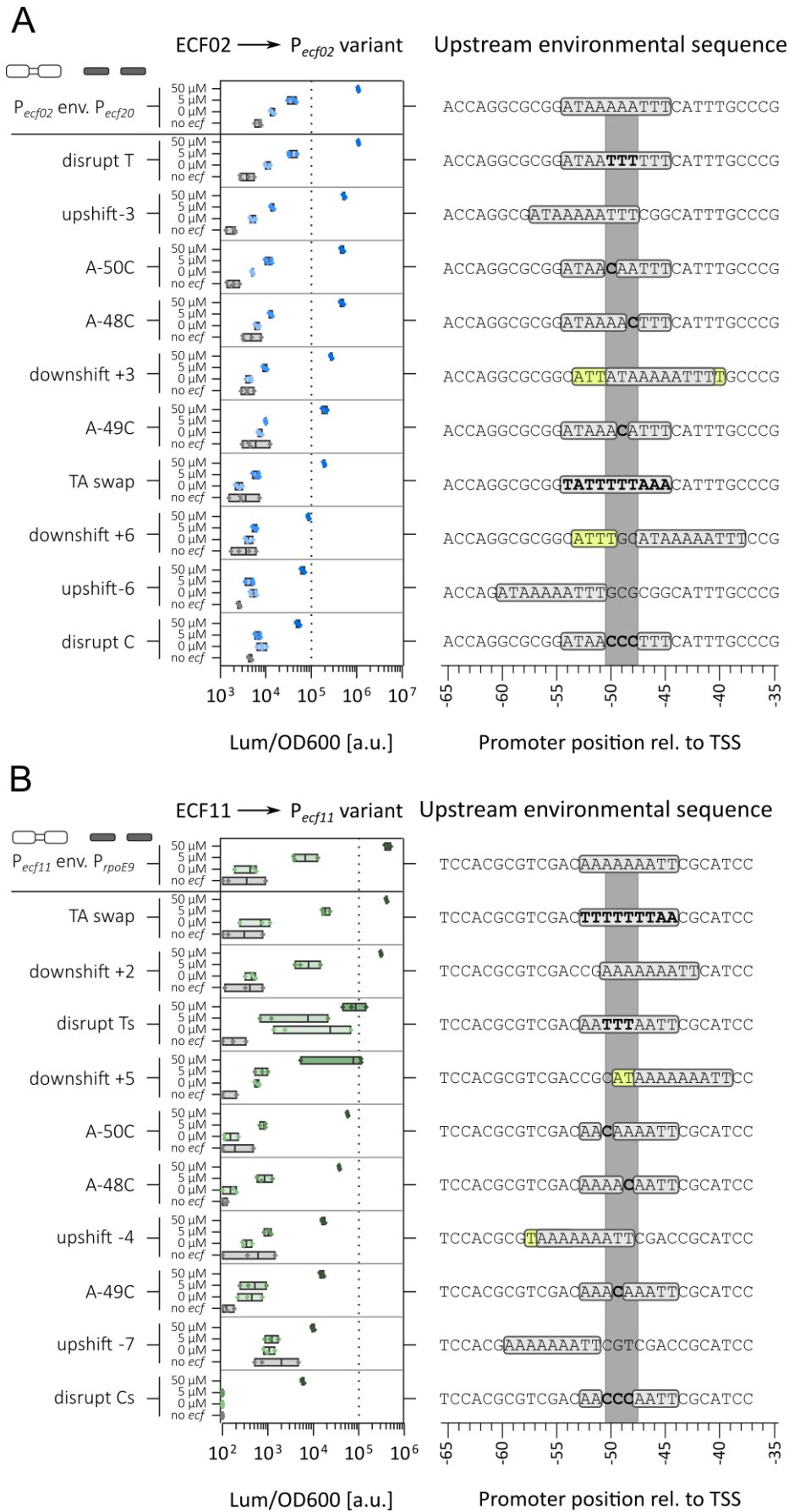

**Figure S7: Mutagenesis of A<sub>n</sub>T motif in two different hybrid promoters using two different environmental sequences.** Mutation analysis for the $P_{ecf02}$ core promoter with the environment of $P_{ecf20}$ (A) and the $P_{ecf11}$ core promoter with the environment of $P_{rpoE9}$ (B). Two plasmid-based ECF switches were measured in *S. meliloti* $\Delta ecf/\Delta anti-\sigma$ in MOPS-buffered

minimal medium and OD600 and luminescence were capture 9 h after addition of salicylate (0 μM, 5 μM, 50 μM). Each switch was controlled by a strain that carried an empty plasmid instead of the input plasmid ("no *ecf*" control). The promoters in both switches were subject to the following mutations: (i) Changing the common A<sub>n</sub>T motif bases -50, -49, and -48 to thymines (T-disruption) or cytosines (C-disruption). (ii) Single point mutations of each base common to all A<sub>n</sub>T motifs found to cytosines (A-50C, A-49C, and A-48C). (iii) Exchanging all adenines in the entire A<sub>n</sub>T motif to thymines and vice versa (TA-swap). (iv) Repositioning of the A<sub>n</sub>T motif upstream or downstream to either directly outside the common A<sub>n</sub>T motif positions or such that the A<sub>n</sub>T motif just covered the common positions. Each variant of the respective promoter is shown in the right. The grey bar marks the common A<sub>n</sub>T motif bases, light grey boxes show the A<sub>n</sub>T motif and its position. Bases that were altered in the A<sub>n</sub>T motif are indicated in bold font. If repositioning of the A<sub>n</sub>T motif introduced a longer A/T stretch that was not part of the original A<sub>n</sub>T motif, these bases are highlighted in yellow boxes. Data represents at least three biological replicates.

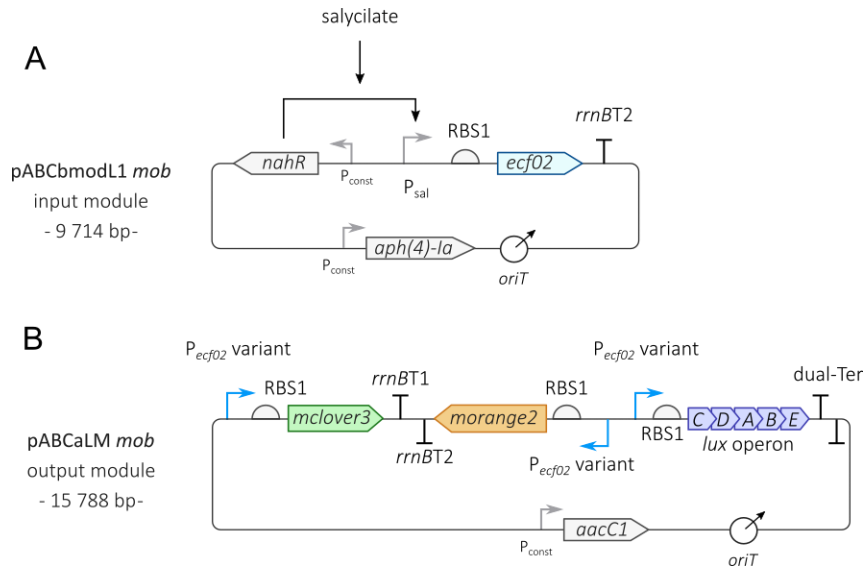

**Figure S8: Implementation details of the ECF02-dependent 3-regulon switch.**

The 3-regulon was divided in two modules each on a separate single-copy number plasmid: The input module (**A**) carried the *ecf02* gene under the control of a salicylate-inducible promoter on a pABCbmodL1 *mob* plasmid that conferred resistance to hygromycin-B. This is the same input plasmid used for 1-regulon switches. The output module (**B**) carried three different reporters (the fluorescent genes *mclover3* and *morange2* as well as the luminescence cassette *luxCDABE*) under the transcriptional control of ECF02-dependent promoter variants. Each individual transcription unit consisted of a common ribosome binding site (RBS) as part of the 5'-UTR and different transcriptional terminators.

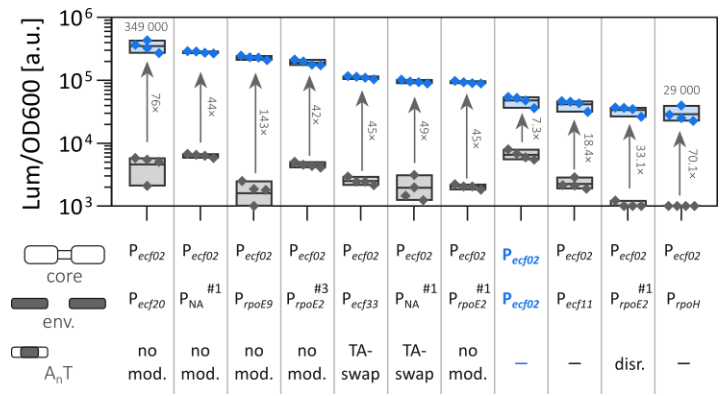

**Figure S9: Activity of ECF02-dependent two-plasmid based switches with a common set of environmental hybrid promoters.**

Luminescence output of *S. meliloti* $\Delta ecf/anti-\sigma$ carrying ECF02-dependent two plasmid switches with different promoters in MOPS-buffered minimal medium 9 h after the addition of 0 μM (grey symbols) or 50 μM (blue symbols) salicylate. Numbers on top of the boxes indicate the average relative luminescence of the on-states of the strongest and weakest switch. Numbers on the arrows show average fold change between on- and off-state of each switch. Descriptions show the core motif, promoter environments used, and presence of or modifications to the A<sub>n</sub>T motif in the environment (abb.: "no mod." – no modification; "TA-swap" – all adenine and thymine bases swapped; "disr." – disruption of the motif with C bases; "—" – no A<sub>n</sub>T motif present in the environment). Bold blue label indicates the native P<sub>ecf02</sub> promoter. Data based on four biological replicates

Figure S10: An artificial ECF02 regulon consisting of three regulated genes.

Luminescence output of S. meliloti Δecf/Δanti-σ carrying ECF02-dependent 1-regulon (A-C) and 3-regulon swichtes (D-F) in MOPS-buffered minimal medium in the presence or absence or 50 µM salicylate. Relative mClover3 and mOrange2 fluorescence was captured 23 h and relative luminescence was captured 6 h post induction. The Pecf02 promoter variant that was used to drive the expression of respective reporter(s) is indicated by the colored boxes with symbols (black (+), grey (~), and white (-), respectively) beneath the plots where “X” represents the absence of a reporter in the 1-regulons. This figure represents the same cultivation as Fig. 6.

**-RNAP $\alpha$ -CTD positioning may vary with different ECF-type sigma factors**

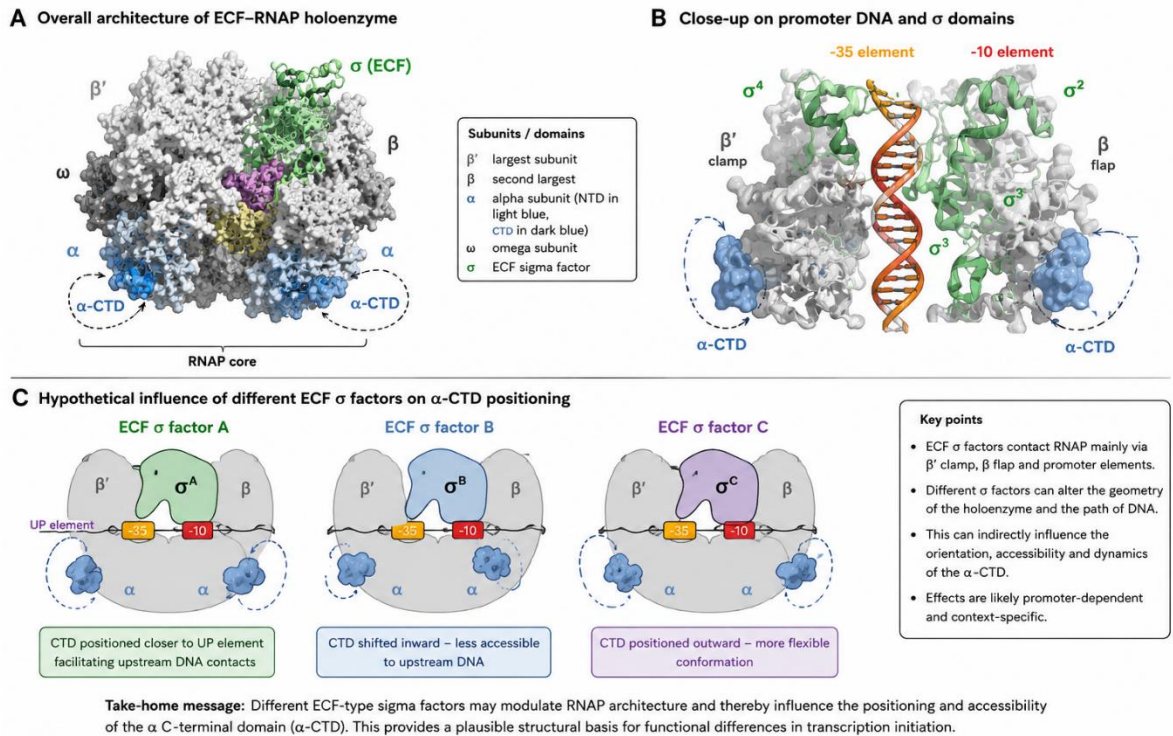

**Figure S11. Hypothetical influence of ECF $\sigma$ factors on RNAP α-CTD positioning and promoter engagement.**

**(A)** Overall architecture of a bacterial ECF–RNAP holoenzyme based on the available *E. coli* RpoE-containing complex structure (PDB: 6JBQ). The RNA polymerase (RNAP) core enzyme is shown in grey, the ECF $\sigma$ factor in green, and the α-subunit C-terminal domains (α-CTDs) in blue. Due to their intrinsic flexibility, α-CTDs are thought to sample multiple conformational states relative to the RNAP core and upstream promoter DNA. **(B)** Close-up view of promoter engagement by ECF domains. Conserved sigma domains interact primarily with the $-35$ and $-10$ promoter elements, while the α-CTDs may contact upstream DNA regions or UP-like elements depending on promoter architecture and holoenzyme conformation. Dashed arrows indicate the potential conformational mobility of the α-CTDs. **(C)** Conceptual model illustrating how distinct ECFs may indirectly alter α-CTD positioning and accessibility through changes in RNAP holoenzyme geometry, promoter DNA trajectory, or σ-dependent promoter interactions. Different conformational states could influence the ability of α-CTDs to engage upstream DNA regions, thereby modulating transcription initiation efficiency in a promoter-dependent manner.
